# Rapid evolution of the fine-scale recombination landscape in wild house mouse (*Mus musculus*) populations

**DOI:** 10.1101/2022.06.08.495013

**Authors:** Lydia K. Wooldridge, Beth L. Dumont

**Affiliations:** The Jackson Laboratory, 600 Main Street, Bar Harbor ME 04609; Tufts University, Graduate School of Biomedical Sciences, 136 Harrison Ave, Boston MA 02111

## Abstract

Meiotic recombination is an important evolutionary force and essential meiotic process. In many species, recombination events concentrate into “hotspots” defined by the site-specific binding of PRMD9. Rapid evolution of PRDM9’s zinc finger DNA-binding array leads to remarkably abrupt shifts in the genomic distribution of hotspots between species, but the question of how *Prdm9* allelic variation shapes the landscape of recombination between populations remains less well understood. Wild house mice (*Mus musculus*) harbor exceptional *Prdm9* diversity, with >100 alleles identified to date, and pose a particularly powerful system for addressing this open question. We employed a coalescent-based approach to construct fine-scale, sex-averaged recombination maps from contemporary patterns of linkage disequilibrium in nine geographically isolated wild house mouse populations, including multiple populations from each of three subspecies. Comparing maps between wild mouse populations and subspecies reveals several themes. First, we report weak fine- and broad-scale recombination map conservation across subspecies and populations, with genetic divergence offering no clear prediction for recombination map divergence. Second, most hotspots are unique to one population, an outcome consistent with minimal sharing of *Prdm9* alleles between surveyed populations. Finally, by contrasting aggregate hotspot activity on the X versus autosomes, we uncover evidence for population-specific differences in the degree and direction of sex-dimorphism for recombination. Overall, our findings illuminate the variability of both the broad- and fine-scale recombination landscape in *Mus musculus* and underscore the functional impact of *Prdm9* allelic variation in wild mouse populations.

## Introduction

Recombination is both an important evolutionary mechanism for generating genetic diversity and an essential meiotic process. At least one crossover per chromosome is required for proper synapsis and segregation of homologous chromosomes during the first meiotic division, with too few, too many, or improperly positioned crossovers resulting in the production of aneuploid gametes (Hassold and Hunt 2001; Ferguson et al. 2007). Despite its critical importance for faithful genome transmission, recombination rates show extreme variation between species, between populations, and among individuals. Recent studies have demonstrated that a significant proportion of this variation is under genetic control and have also identified environmental variables that contribute to recombination rate plasticity (Hunt et al. 2003; Hunter et al. 2016; Henderson and Bomblies 2021; Belmonte-Tebar et al. 2022). However, the evolutionary forces that shape recombination rate variation in nature remain largely enigmatic.

In many mammals, including mice and humans, recombination is sexually dimorphic. For example, in humans, females have higher average crossover counts than males, whereas males exhibit an enrichment of crossovers near the telomeric ends of chromosomes (Kong et al. 2002; Paigen et al. 2008). These sex differences in recombination rate and distribution are also observed in most inbred laboratory strains of house mice (Dumont and Payseur 2011a). Intriguingly, however, a select number of inbred strains have recently been identified that exhibit a reversal in the usual direction of the sex dimorphism: In strains PWD/PhJ and MSM/MsJ, females have lower recombination rates than males (Peterson and Payseur 2021). These discordant findings suggest that the directionality of the sex dimorphism for recombination rate can also evolve rapidly, potentially driven by sex-specific selection for distinct recombination rates in male and female meiosis (Dumont and Payseur 2011b). However, as no studies have yet surveyed male and female recombination rates in outbred wild mouse populations, it remains unclear whether the higher male recombination rates observed in strains like PWD/PhJ and MSM/MsJ are mere artifacts of inbreeding.

In addition to varying between genomes, recombination rates are also heterogeneous within genomes. On the scale of megabases, recombination rates tend to be elevated near telomeres and suppressed in heterochromatic centromeric regions (Nachman and Churchill 1996; Kong et al. 2002; Jensen-Seaman et al. 2004). On broad scales, recombination rates also co-vary with respect to numerous genomic features, including gene density, GC content, and repetitive DNA (Kong et al. 2002; Jensen-Seaman et al. 2004; Buard and de Massy 2007; Brick et al. 2012). At finer physical scales, the recombination landscape is dominated by the positioning of small 1-5 kb recombination ‘hotspots’. Virtually all recombination events concentrate into hotspots, meaning that most of the genome is recombinationally inert and never participates in recombination (McVean et al. 2004).

In many mammalian species, the location of recombination hotspots is defined by the zinc finger protein PRDM9 (Baudat et al. 2010). PRDM9 localizes to specific DNA binding sequences recognized by its zinc finger domain. Once bound, PRDM9 trimethylates local histones at both H3K4 and H3K36 (Powers et al. 2016). This epigenetic signature recruits the double-strand break machinery to the site to initiate a cascade of DNA repair events that culminate in the formation of crossovers or non-crossover gene conversion events. Comparative genomic investigations have revealed that the zinc finger array of *Prdm9* evolves rapidly, leading to abrupt changes in the suite of PRDM9 binding sequences across the genome and concomitant shifts in the fine-scale genomic distribution of recombination hotspots (Oliver et al. 2009; Baker et al. 2017). As a result, recombination hotspots exhibit minimal conservation between species (Stevison et al. 2016), although there are appreciable levels of hotspot sharing between human populations (Spence and Song 2019).

While recent investigations in laboratory mice have shed light on the molecular mechanisms of PRDM9 action and defined strain differences in PRDM9-dependent recombination hotspot distribution, the question of how *Prdm9* allelic variation shapes the landscape of recombination in wild populations remains less well understood (Brick et al. 2012; Powers et al. 2016; Grey et al. 2018). More than 100 *Prdm9* alleles have been characterized in wild mice to date, with most alleles restricted to single populations and few shared between subspecies (Buard et al. 2014; Vara et al. 2019). These aspects of the population genomic distribution of *Prdm9* allelic variation predict substantial population and subspecies level diversity in the fine-scale distribution of recombination hotspots.

Fine-scale variation in recombination – and in particular the location of hotspots – within a population can exert profound effects on population evolution and diversity. For one, recombination influences haplotype diversity within populations by shuffling alleles between homologous chromosomes. In addition, by breaking down associations between high fitness alleles and linked deleterious variants, recombination can reduce selective interference and expedite the fixation of adaptive alleles (Crow and Kimura 1965; Maynard Smith 1971). All else being equal, an adaptive variant that arises in a high recombination rate region is expected to reside on a shorter haplotype and encounter less selective interference than a high fitness allele that emerges in a recombination coldspot (Hey 2004). Conversely, the extent of the reduction in flanking diversity accompanying selection against a deleterious allele depends on the local recombination rate and the precise positioning of hotspots (Charlesworth et al. 1993). Thus, knowledge of the fine-scale recombination landscape is essential for a holistic interpretation of standing patterns of population diversity.

Multiple approaches for measuring fine-scale recombination rates have been developed, each offering distinct strengths and weaknesses. Bulk genotyping of sperm from single individuals can reveal the frequency of recombinant haplotypes at targeted loci (Jeffreys et al. 2001; Jeffreys et al. 2004). While this approach is highly sensitive and can be readily scaled to multiple samples, it cannot be used to comprehensively interrogate fine-scale recombination rates genome-wide, nor can it be adapted to probe female recombination rates. Single-cell technologies have recently been used to ascertain the recombination landscape in sperm and oocytes from single individuals (Wang et al. 2012; Hou et al. 2013; Ottolini et al. 2015; Dréau et al. 2019; Bell et al. 2020). However, these methods remain prohibitively expensive to apply to large numbers of samples, barring their application at the population scale. Bulk sequencing of DNA fragments bound to recombination-associated proteins provides a third strategy for surveying the fine-scale landscape of meiotic recombination (Smagulova et al. 2011; Khil et al. 2012). However, this approach is similarly cost- and time-prohibitive at scale.

A fourth approach for defining the fine-scale recombination landscape relies on population genomic analyses of whole genome sequences or dense SNP data from population samples. This approach is premised on the insight that the level of linkage disequilibrium (LD) between two loci in a given population offers a read-out of the historical rate of recombination between those sites (McVean et al. 2004). Thus, by surveying patterns of genetic variation in contemporary populations, one can obtain estimates of the population-scaled recombination rate, rho (*ρ*), between every pair of segregating sites in the genome, yielding the finest possible recombination map resolution. These estimates reflect the cumulative recombination activity of all individuals in the population and over the history of the population, and therefore provide a time- and sex-averaged portrait of fine-scale recombination activity. However, as the X chromosome only engages in recombination in the female germline, contrasts between recombination rates on the X and autosomes, which recombine in both sexes, may be especially informative about sex differences in meiotic recombination.

Here, we use this latter approach to generate broad- and fine-scale genome-wide recombination maps from whole genome sequences of wild-caught mice from nine geographically-isolated locations (Davies 2015; Harr et al. 2016). These surveyed populations include multiple populations from each of the three principal house mouse subspecies: *M. m. domesticus* (Germany, Iran, 2 populations from France), *M. m. musculus* (Kazakhstan, Afghanistan, Czech Republic), and *M. m. castaneus* (India, Taiwan). We then use these maps to address several outstanding questions. First, do levels of broad-scale recombination rate divergence scale with population and subspecies divergence? Second, what is the extent of fine-scale recombination rate variation among wild house mouse populations and subspecies? Third, is there evidence for population differences in the polarity of sex-dimorphism for recombination rate? Taken together, our findings provide a window into the evolutionary history of fine- and broad-scale recombination rates in wild house mice, extending insights gleaned from inbred mouse strains and exposing the functional consequences of the exceptional *Prdm9* diversity in *M. musculus*.

## Results

### Sequencing data summary and switch-error rates

We utilized publicly available whole-genome sequences from wild-caught mice from nine geographic locations to estimate population-specific recombination maps and hotspot locations (Davies 2015; Harr et al. 2016). We refer to the nine populations as: mAfghanistan, mCzechia, mKazakhstan, dIran, dGermany, dFrance_1, dFrance_2, cTaiwan, and cIndia, with the leading letter denoting the primary subspecies designation of each population (m: *musculus*; d: *domesticus*; c: *castaneus*). After quality control filtering, a range of 7,908,349 (mAfghanistan) to 40,890,538 (cIndia) SNPs were identified per population (mean 17,427,800 SNPs), corresponding to approximately one SNP per ∼60-300 bp, on average (Table 1).

**Table 1:** Whole Genome Sequence Data Summary

| Sub-Species | Population | Source | # Samples | # Males | # SNPs | SNP density (bp/SNP) | Switch-error Rate (%) |
| --- | --- | --- | --- | --- | --- | --- | --- |
| Castaneus | India | <sup>a</sup> | 10 | 3 | 40,890,538 | 60 | 0.24 |
| Castaneus | Taiwan | <sup>b</sup> | 20 | 1 | 25,549,656 | 96 | na |
| Domesticus | Iran | <sup>a</sup> | 8 | 8 | 17,877,283 | 138 | 0.039 |
| Domesticus | Germany | <sup>a</sup> | 11 | 9 | 11,930,888 | 206 | 0.056 |
| Domesticus | France_1 | <sup>a</sup> | 8 | 8 | 11,108,085 | 222 | 0.081 |
| Domesticus | France_2 | <sup>b</sup> | 20 | 10 | 14,120,193 | 174 | 0.18 |
| Musculus | Afghanistan | <sup>a</sup> | 6 | 5 | 7,908,349 | 311 | 0.79 |
| Musculus | Czechia | <sup>a</sup> | 8 | 2 | 10,208,203 | 241 | na |
| Musculus | Kazakhstan | <sup>a</sup> | 8 | 4 | 10,937,288 | 225 | 0.34 |

<sup>a</sup> (Harr et al. 2016)
<sup>b</sup> (Davies 2015)

SNPs were computationally phased into haplotypes. Errors in haplotype inference will masquerade as recombinants and may artificially inflate estimates of the population scaled recombination rate, *ρ*. To assess the incidence of such haplotype ‘switch-errors’ in our data, we randomly paired phase-known X chromosome haplotypes in males to generate pseudofemales that were used to directly benchmark the switch error rate in most populations (see Methods). On average across populations, the switch-error rate is 0.25%, ranging from 0.04 to 0.79% among populations (Table 1). These error rates are comparable to or lower than those reported in prior investigations (Booker et al. 2017; Shanfelter et al. 2019).

### Population-scaled recombination rates reflect general features of house mouse demographic history

Across the nine surveyed populations, the mean *ρ*/bp estimate for all chromosomes ranged ∼12-fold (Figure 1), from a low of 0.0010037 *ρ*/bp (dGermany) to a high of 0.01233 *ρ*/bp (cIndia). Autosome means ranged from 0.001032 (dGermany) to 0.012766 (cIndia) *ρ*/bp, while the X mean ranged from 0.00114 (dGermany) to 0.0043 (mKazakhstan) *ρ*/bp. The mean *ρ*/bp for individual chromosomes from each population is provided in Supp. File 1.

**Figure 1:**
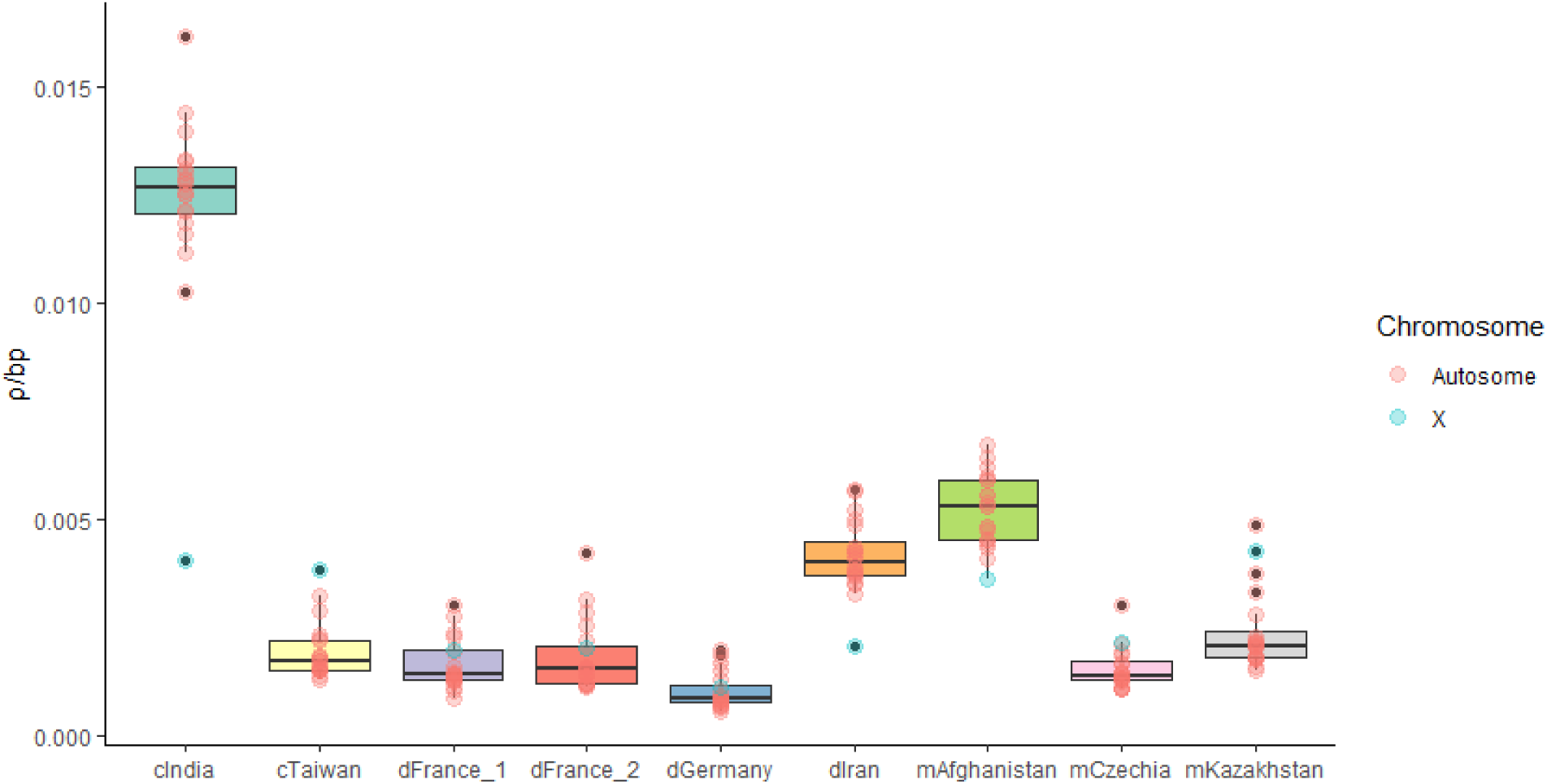
Estimated *ρ*/bp for each chromosome and each population. The mean *ρ* of all 20 chromosomes is summarized as a box-and-whisker plot.

House mice evolved from a common ancestral source population in the Indo-Iranian valley approximately 0.5 million years ago (Boursot et al. 1993). Mean *ρ*/bp estimates were 2-12 times higher in mice collected from regions closest to this ancestral region (India, Iran and Afghanistan) compared to more derived populations. These trends reflect, in large part, the higher historical effective population sizes of the ancestral populations. Given the slight tendency to over-estimate population scaled recombination rates when the true *ρ*/bp is low (<0.002) and when switch-error rates are moderately high (>0.46%) (Booker et al. 2017), estimates for several populations may be weakly inflated (mCzechia, dGermany, dFrance_1, dFrance_2, cTaiwan). Thus, the magnitude of reported population differences in *ρ*/bp is potentially conservative.

### Weak conservation of broad-scale recombination maps across M. musculus populations and subspecies

To compare recombination rates across these nine populations of mice, we first translated recombination rate estimates from *ρ*/bp to cM/Mb units and averaged the resulting rate estimates over 1Mb windows (see Methods). To ensure the robustness of this approach, we compared our broad-scale chromosome-level maps for cIndia to previously generated recombination maps for this population (Booker et al. 2017). Despite differences in methodology and genome builds, concordance between these broad-scale maps is excellent (mean Spearman *ρ* across chromosomes = 0.89; per chromosome range 0.64-0.96; *P* < 0.05; Figure 3D).

We next assessed the similarity of recombination rates in 1Mb windows between each pair of wild mouse populations. Map correlations are expected to decline with genetic divergence (Stevison et al. 2016), and we anticipated that recombination maps would exhibit greater similarity between populations of the same *Mus musculus* subspecies, relative to populations from different subspecies. Average map correlations were 0.4 (Range: 0.31 - 0.52), 0.38 (Range: 0.35 – 0.41), and 0.44 for comparisons within *domesticus*, *musculus*, and *castaneus*, respectively (Spearman’s *ρ*; all comparisons, *P* << 1x10^-10^; Figure 2). However, in contrast to our expectations, the average correlation between recombination maps for inter-subspecies comparisons was of identical magnitude (Spearman’s *ρ* = 0.4, all with *P* << 1x10^-10^; Figure 2). The map comparisons between dIran and cIndia yielded the highest correlation (Spearman’s *ρ* = 0.62, *P* << 1x10^-10^), potentially reflecting the ancestral identity of these populations (Figure 2). Examples of the magnitude of spatial and population variation in broad-scale recombination rates are presented in Figures 3A-C. Correlations for individual chromosome comparisons are presented in Supp. File 2.

**Figure 2:**
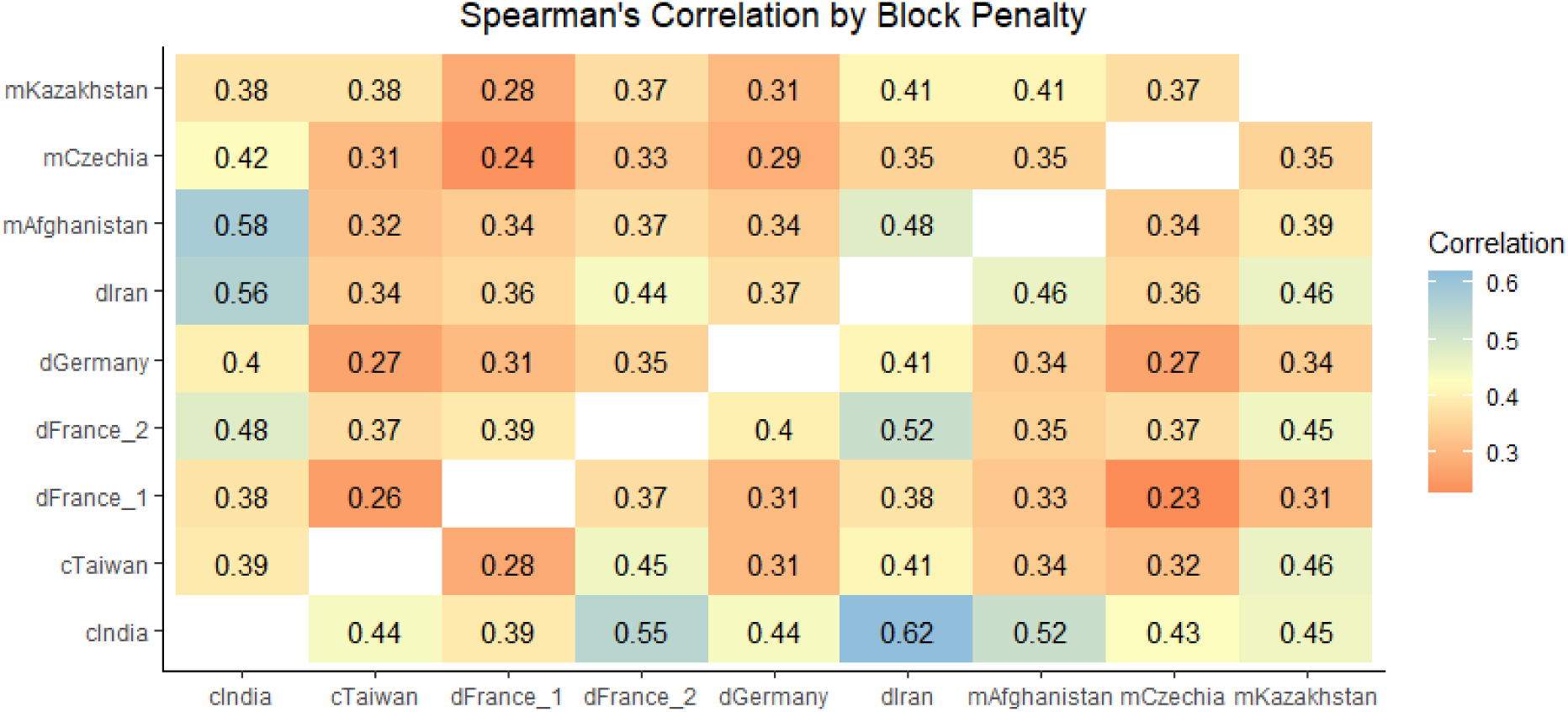
Heat map of Spearman’s rank correlation values for all inter-population comparisons. Both block penalty results are shown, with correlations between maps constructed with a block penalty of 10 presented above the diagonal, and correlations between maps constructed under a block penalty of 100 shown below the diagonal.

**Figure 3:**
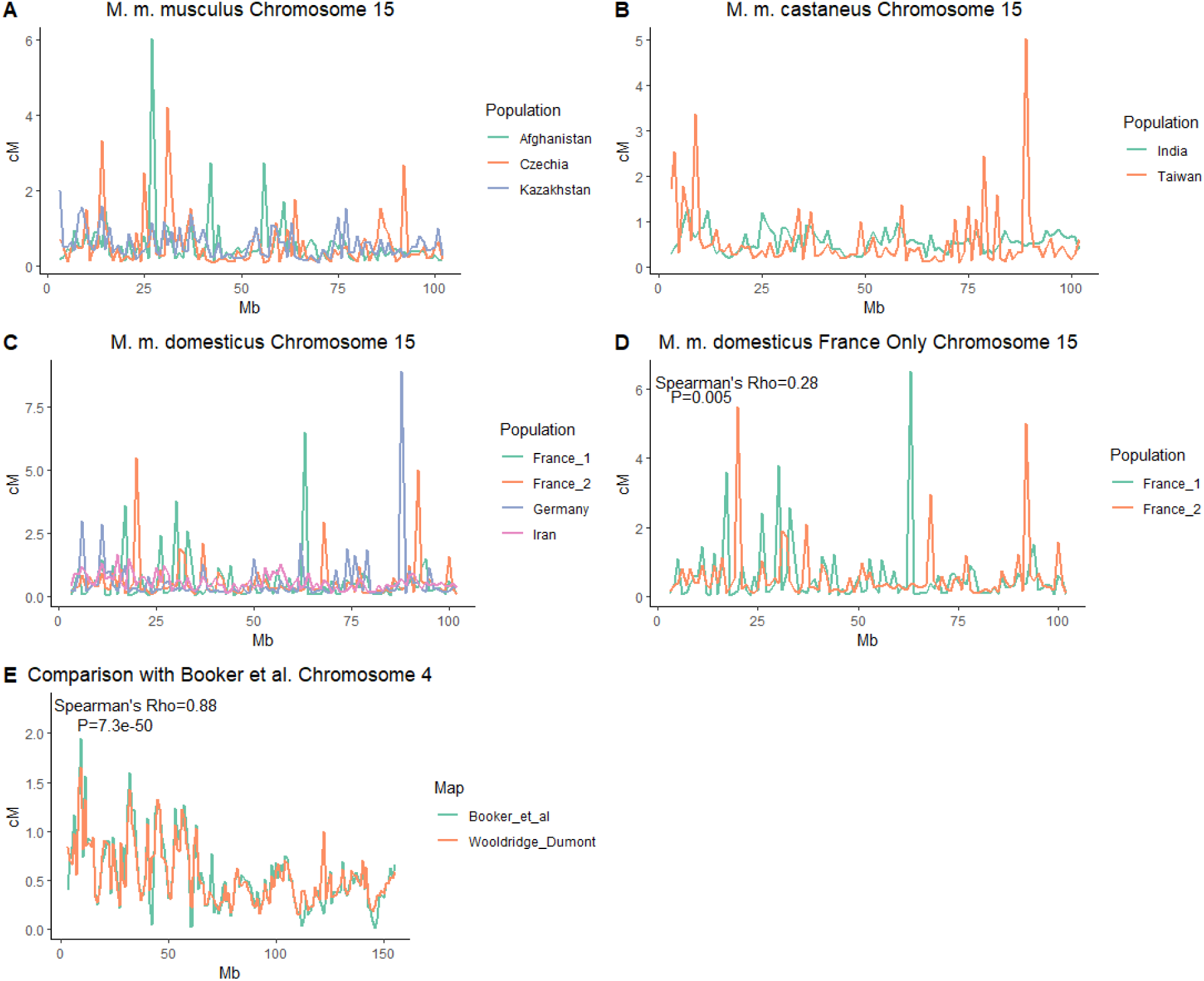
Comparisons of broad-scale recombination maps across *Mus musculus* populations in cM/Mb units. A) Chromosome 15 for all *M. m. musculus* populations, B) chromosome 15 for all *M. m. castaneus* populations, C) chromosome 15 for all *M. m. domesticus* populations, D) chromosome 15 for only the two *M. m. domesticus* populations from France, and E) a comparison of our chromosome 4 cIndia map with the map generated from the same data by Booker et al. (Booker et al. 2017). Chromosome 4 was selected for E because the Spearman’s correlation for this chromosome is similar to the mean correlation for all chromosomes.

To determine if these inter-population correlations in broad-scale recombination rates are higher than expected by chance, we randomly permuted *ρ*/bp estimates in 1 Mb windows across the whole genome and re-assessed correlations between populations. The mean permutation-based correlation per population comparison ranged from -0.005 to 0.003. In 100 permutation replicates per comparison, correlations never exceeded the values recovered from the actual maps (*P* < 0.01). In summary, the strength of observed correlations between *Mus musculus* broad scale maps do not scale with population and subspecies divergence, but nonetheless remain significantly higher than expected by chance.

### Fine-scale recombination maps are weakly conserved across populations

We next compared fine-scale recombination maps between populations. Due to the rapid evolutionary turnover of recombination hotspots, we expected to recover reduced correlations in fine-scale map comparisons relative to comparisons between the broad-scale maps. In line with these predictions, most inter-population fine-scale map comparisons exhibited weaker correlation than the corresponding broad-scale map comparisons (28/36 comparisons), although the difference in correlation magnitude is modest (Figure 2). Correlation magnitudes are similar for all within subspecies fine-scale map comparisons (Spearman’s *ρ* = 0.37, 0.36, and 0.39 for *domesticus*, *musculus* and *castaneus*, respectively; all *P* << 1x10^-10^), and comparable to the strength of observed correlations for inter-subspecies fine-scale map comparisons (average Spearman’s *ρ* = 0.37; all *P* << 1x10^-10^).

### Hotspot identification

We used two approaches to comprehensively identify historical recombination hotspots in each surveyed wild population (see Methods). Briefly, the ‘sliding window’ hotspot method, which has been used in prior analyses (Booker et al. 2017; Shanfelter et al. 2019), tests whether the *ρ* estimate of every genomic window of a pre-defined, fixed length is significantly greater than the population-scaled recombination rate of the flanking regions. If so, such regions are identified as hotspots. This approach fails to fully leverage the high density of SNPs in whole genome sequencing datasets and may over-estimate hotspot size. To circumvent these potential shortcomings, we developed a second approach (the ‘filtering’ method) which identifies hotspots as inter-SNP intervals with *ρ*/bp estimates significantly greater than the chromosome-wide mean *ρ*/bp. Adjacent intervals meeting these criteria are merged into a single candidate hotspot, with hotspots defined by >2 SNPs and <5 kb in length retained.

Using the sliding window method, we identified a total of 225,605 hotspots across all populations of mice, with a mean of 25,067 hotspots per population (Table 2). Using the filtering method, we identified 214,717 hotspots, with an average of 23,857 hotspots per population (see Supp. File 3). The number of putative hotspots identified by both approaches are similar to previous reports for *Mus musculus* (Brunschwig et al. 2012; Smagulova et al. 2016; Booker et al. 2017).

**Table 2:** Hotspot Counts and General Characteristics

| Population | Sliding Window Hotspots |  |  | Unique Sliding Window Hotspots |  | Filtered Hotspots |  |  | Unique Filtered Hotspots |  |
| --- | --- | --- | --- | --- | --- | --- | --- | --- | --- | --- |
| | Number | Mean Length (bp) | Mean $\rho$/bp | Mean Length (bp) | Mean $\rho$/bp | Number | Mean Length (bp) | Mean $\rho$/bp | Mean Length (bp) | Mean $\rho$/bp |
| mAfghanistan | 27,849 | 2,222 | 0.33 | 2,232 | 0.23 | 19,135 | 1,158 | 741 | 0.74 | 0.45 |
| mCzechia | 20,497 | 2,067 | 0.14 | 2,066 | 0.08 | 12,117 | 756 | 420 | 0.36 | 0.11 |
| mKazakhstan | 25,182 | 1,759 | 0.07 | 1,729 | 0.04 | 17,463 | 475 | 292 | 0.33 | 0.16 |
| dIran | 34,407 | 1,723 | 0.21 | 1,624 | 0.10 | 32,419 | 579 | 366 | 0.52 | 0.23 |
| dGermany | 20,684 | 1,881 | 0.05 | 1,859 | 0.03 | 14,711 | 682 | 422 | 0.14 | 0.06 |
| dFrance_1 | 22,772 | 1,952 | 0.13 | 1,921 | 0.07 | 13,897 | 686 | 389 | 0.32 | 0.12 |
| dFrance_2 | 23,749 | 1,886 | 0.07 | 1,796 | 0.03 | 19,819 | 705 | 447 | 0.16 | 0.07 |
| cIndia | 28,190 | 1,502 | 0.31 | 1,402 | 0.13 | 56,134 | 368 | 0.65 | 257 | 0.46 |
| cTaiwan | 22,275 | 1,677 | 0.08 | 1,563 | 0.03 | 29,022 | 472 | 0.15 | 340 | 0.08 |

We evaluated the extent of hotspot overlap between these two hotspot calling methods and defined key features of hotspots identified by these two approaches (Table 2). Of the 225,605 hotspots identified by the sliding window approach, 115,978 (51.4%) were not called using our filtering method (109,627 hotspots (48.6%) are shared between the two methods). Of the 214,717 filtered hotspots, 100,061 (46.6%) are uniquely ascertained by this approach (114,656 filtered hotspots (53.4%) were also identified by the sliding window method). Of the 114,656 filtered hotspots that overlap with a sliding window hotspot, 9,808 (8.6%) did not fully overlap the sliding window hotspot. Conversely, all sliding window hotspots overlapping a filtered hotspot showed complete overlap with the filtered hotspot.

Mean hotspot length was 1,852 bp for all sliding window hotspots versus 654 bp for filtered hotspots. When hotspots detected by both methods were removed, the mean length of sliding window hotspots was reduced to 1,799 bp and to 408 bp for filtered hotspots. The mean recombination rate for the sliding window hotspots (in *ρ*/bp) was 0.15, but only 0.08 for hotspots uniquely called by this method. Hotspots called by the filtering method were considerably ‘hotter’, and averaged 0.37 (*ρ*/bp) for the entire dataset, and 0.19 (*ρ*/bp) for hotspots unique to this method. This distinction is likely due to the smaller size of filtered hotspots, which excludes the dampening impact of recombinationally inert flanking sequences.

Because many hotspots were unique to a given calling method, we merged the two datasets to obtain a comprehensive hotspot dataset. Overlapping hotspots (minimum 1 bp overlap) were merged to create a single larger hotspot. Overall, we identified 325,604 putative hotspots among the nine mouse populations. On average for a given population, these combined hotspots covered approximately 1.85% of the genome (in terms of bp), reinforcing the point that recombination activity is concentrated into a small proportion of the genome.

### Hotspots are under-represented in repetitive elements

Repetitive elements comprise a large fraction of the mouse genome and are broadly associated with reduced recombination rates (Jensen-Seaman et al. 2004). To confirm this genomic association, we compared the relative density of different classes of repetitive elements in hotspots versus coldspots. Regardless of the hotspot calling method, almost all repetitive elements were proportionately more abundant (Fisher’s Exact test; *P* < 0.05) in coldspots, with few exceptions (See Supp. File 4). In cIndia, B4 and MIR repeats were more prevalent (*P* < 0.05) in sliding window hotspots than coldspots.

We next determined the percentage of each hotspot or coldspot overlapping annotated repetitive elements in each population. For most populations, sliding window hotspots exhibited lower percent overlap with repeats than coldspots (overlap quantified as the percentage of overlapping base pairs; Welch’s T-test; *P*<0.05). In contrast, filtered hotspots displayed the opposite trend, with greater overlap between hotspots and repeats than between coldspots and repeats (See Supp. File 5).

Filtered hotspots exhibit a lower overall density of repetitive elements than coldspots, but increased repeat composition of hotspots. We asked whether these perplexing and seemingly contradictory observations might be attributable to the shorter length of filtered hotspots compared to both sliding window hotspots and coldspots. Length significantly predicted repeat percentage for 20/27 possible comparisons (9 populations x 3 ‘spot’ types; Beta regression; *P* < 0.05), but offered negligible predictive power in these cases (*R*^2^ < 1%; Figure 4). We conclude that there is no meaningful relationship between hotspot length and fractional repeat composition. Instead, we speculate that the unexpected relationship between filtered hotspots and fractional repeat composition may be driven by the small fraction of hotspots that are fully embedded in annotated repeats (Figure 4).

**Figure 4:**
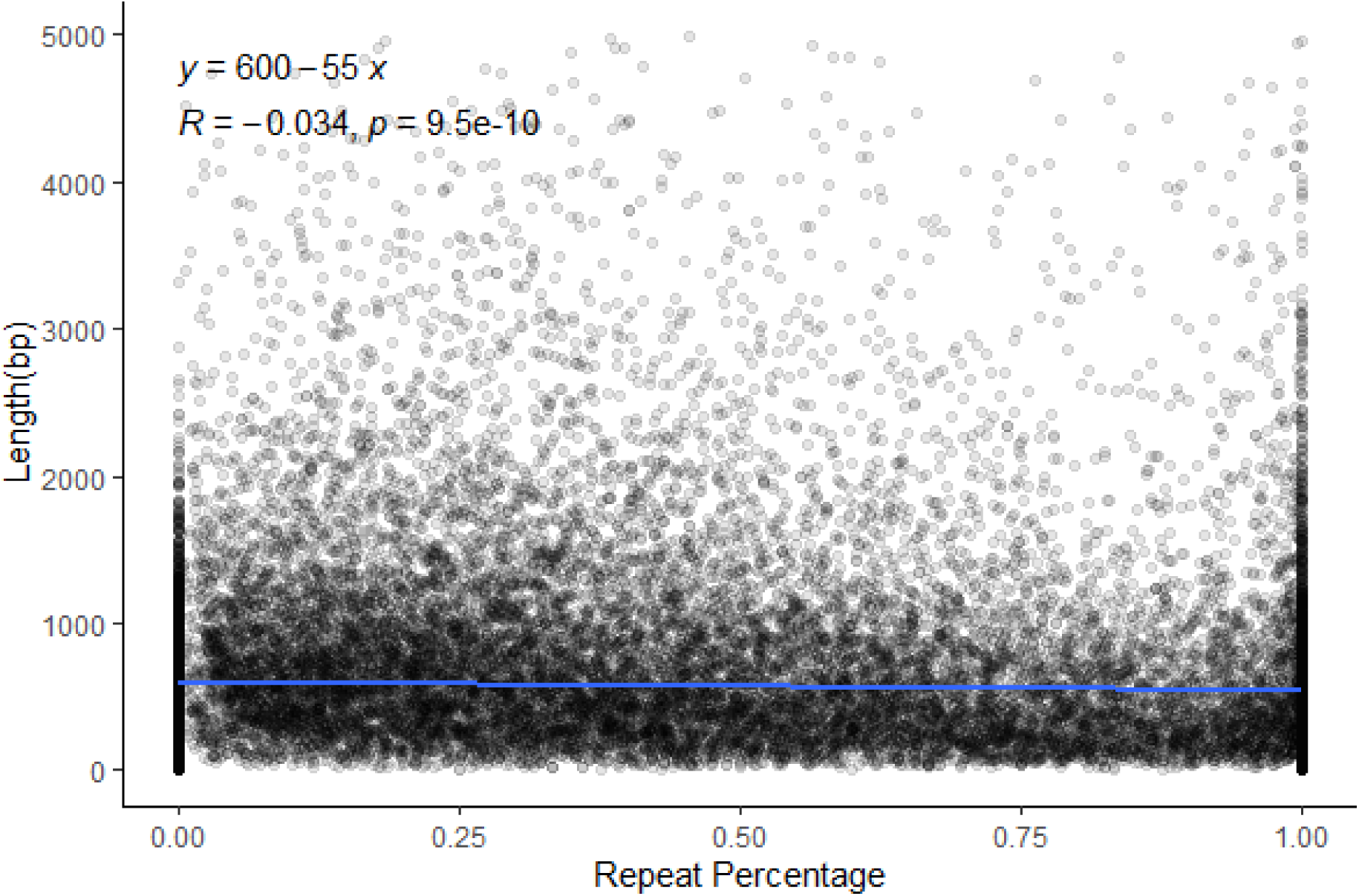
A representative dot plot of the distribution of repeat percentage of filtered hotspots by length for dIran.

### Hotspots differ in proximity to TSS depending on method of ascertainment

PRDM9 sequesters recombination events away from functional genomic elements, shielding them from the potential mutagenic effects of recombination (Brick et al. 2012; Arbeithuber et al. 2015). We therefore predicted that few hotspots should overlap or fall near transcription start sites (TSS). On average, 6% (2.7%) of the sliding window (filtered) hotspots overlapped a TSS. To determine how much overlap could be expected by chance, we simulated ‘randomspots’ across the genome, matching the number and length of observed hotspots. These simulations were repeated 100 times, and the percentage of randomspots overlapping TSS in each simulation run was recorded. In four of the nine populations (mAfghanistan, mCzechia, dGermany and dFrance_1), we observed greater overlap between sliding window hotspots and TSS than expected by chance (Chi-square test; *P* < 0.05). The remaining 5 populations showed no hotspot enrichment with TSSs (*P* > 0.05). A similar pattern was found for the filtered hotspots and their size-matched randomspots: In five of the nine populations (mAfghanistan, mCzechia, dGermany, dFrance_1 and dFrance_2), the observed overlap between filtered hotspots and TSS was significantly greater than expected by chance (*P* < 0.05).

Hotspots ascertained by the two methods differed in their proximity to TSSs. Filtered hotspots from all nine populations were located farther from TSSs than expected by chance (T-test; difference from expectation ranged from +13kb to +73kb, depending on population; *P* << 1x10^-10^). Unexpectedly, for all populations except mKazakhstan, the sliding window hotspots were closer to TSSs than expected (difference ranged from -8kb to -0.7kb across populations; *P* < 0.01). The mKazakhstan sliding window hotspots were only slightly farther from TSS (mean difference: +0.6kb; *P* < 0.01).

Although recombination hotspots are expected to be directed away from TSSs in house mice (Brick et al. 2012), we report an enrichment of hotspots overlapping TSS in 4 to 5 of these 9 wild mouse populations, depending on hotspot calling method. Additionally, the majority of hotspots called by the sliding window method were closer to TSS than expected by chance. These results contradict previous investigations using direct, empirical approaches for detecting meiotic DSBs, the precursors to recombination (Brick et al. 2012). Methodological differences and inevitable false positive hotspots in our dataset (see below) may account for these discrepancies.

Regardless, we note that the proportion of hotspots that overlap TSS in wild house mice is significantly lower than the 20-30% observed in species that lack *Prdm9*-mediated hotspots (Auton et al. 2013; Singhal et al. 2015; Kawakami et al. 2017).

### Assessing hotspot conservation between populations

The rapid evolution of *Prdm9* can lead to wholesale shifts in the fine-scale distribution of recombination hotspots between populations and species. Thus, in species with PRDM9-directed hotspots, geographically isolated populations with distinct *Prdm9* alleles are expected to have relatively few shared hotspots.

We first analyzed how many hotspots were conserved between the surveyed wild *Mus musculus* populations. Using the sliding window hotspots, 2.71 to 14.49% of hotspots overlapped in pairwise population-level comparisons (>50% overlap by bases; mean 4.85%; Figure 5). Comparisons of any combination of the dGermany, dFrance_1 or dFrance_2 mice yielded the highest conservation, potentially reflecting the presence of common *Prdm9* alleles (Buard et al. 2014) due to the recent shared evolutionary history of these populations (Lawal et al. 2021). We observe qualitatively identical trends when examining conservation of filtered hotspots (Range: 1.12 - 10.69% of hotspots overlap by at least 50%; Figure 5) and combined hotspots (Range: 3.26 – 15.0%). However, while overlap between populations was always numerically low, the number of observations is greater than chance expectation (Chi-square test compared with randomspots, *P* << 1x10^-10^). Thus, a minor proportion of hotspots is conserved between populations from the same subspecies.

**Figure 5:**
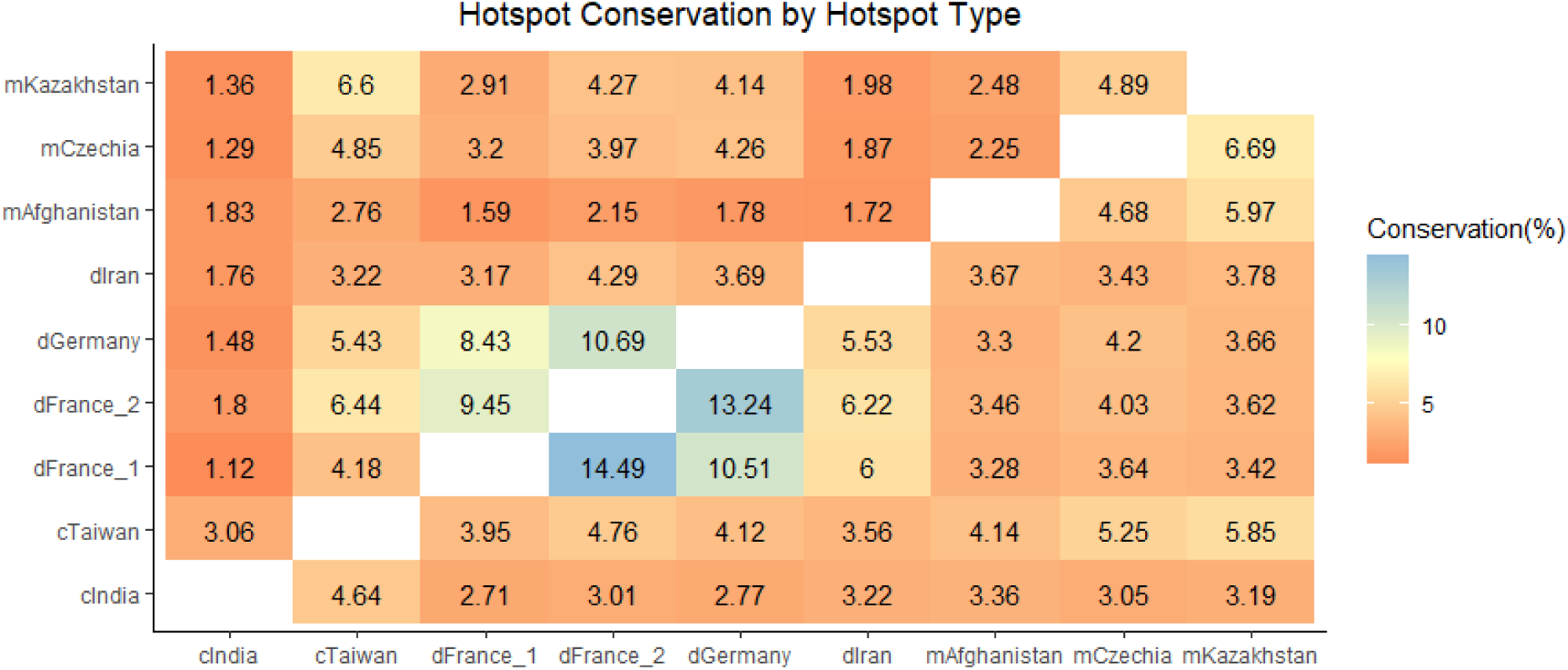
Few hotspots are shared between any two populations. The percentage of hotspots conserved between each population pair is shown as a heat map, with filtered hotspots displayed above the diagonal and sliding window hotspots below the diagonal.

Very few hotspots are conserved (>50% overlap) at the subspecies or species levels. A total of 633 hotspots (combined filtered and sliding window dataset) were shared among all 4 *M. m. domesticus* populations, and 617 were shared across the three *M. m. musculus* populations. The two surveyed *M. m. castaneus* populations share 4,653 hotspots. Only 4 hotspots were common to all populations.

We next used STREME to discover putative PRDM9 sequence motifs enriched in hotspots relative to coldspots for each population. At least one significant motif was detected in 8 of the 9 populations for the filtered hotspots, while only 5 of 9 populations had at least one potential PRDM9 motif detected using the sliding window hotspots (See Supp. Fig. 1). However, some of these putative PRDM9 motifs were present in a small proportion of hotspots (<10%); only 6 of 9 (3 of 9) populations had a motif present in >10% of filtered (sliding window) hotspots. Remarkably, there is no discernable sequence similarity between the most enriched motif in each population, consistent with the historical action of distinct sets of *Prdm9* alleles in each population. Overall, motifs detected from sliding window hotspots were more ambiguous (positions had 2 or more nucleotide possibilities) than those identified from the filtered hotspots (38% ambiguous bases versus 26% ambiguous bases), potentially reflecting the greater length and presence of additional flanking sequence in the former. We further note that each of the identified hotspot-associated motifs is enriched among targets of other known proteins, and therefore the findings from this analysis should be interpreted with caution.

### Hotspot overlap with laboratory strain DSB hotspots

Our analyses reveal minimal hotspot sharing between wild house mouse populations and subspecies. Classical laboratory inbred strains were initially derived from a limited number of wild-caught founder animals and therefore capture a narrow range of the *Prdm9* allelic diversity present in nature (Yang et al. 2011). We sought to determine whether contemporary meiotic double strand break (DSB) positions in diverse inbred strains overlap significantly with the ancestral hotspots discovered in wild mice. We compared our sliding window, filtered, and combined hotspot datasets to the positions of DSB hotspots in 13R, B6, C3H (all *M. m. domesticus*), CAST (*M. m. castaneus*), MOL (*M. m. molossinus*, a hybrid between *castaneus* and *musculus*), and PWD (*M. m. musculus*) inbred mouse strains (Smagulova et al. 2016). Because some overlap is expected by chance, significance was determined by assessing overlap between observed DSB hotspots and simulated randomspots (See Supp. Files 6-7). Overall, we observe an appreciable rate of overlap between DSB hotspots in a given strain and LD-based hotspots ascertained in wild populations from that subspecies (Figure 6). For example, ∼30% of C3H DSB hotspots overlap with LD-hotspots in the two dFrance and dGermany mouse populations of *M. m. domesticus* (4.66 – 6.69% overlap expected by chance). Similarly, we observe ∼20% overlap between CAST DSB hotspots and LD-hotspots in the cTaiwan population (5.32 – 6.17% overlap expected by chance) and ∼30% overlap between PWD DSB hotspots and LD hotspots in the mCzechia population (4.2 – 4.9 % overlap expected by chance). Thus, the *Prdm9* alleles present in modern laboratory mice have left discernable footprints in patterns of LD and the distribution of recombination hotspots in wild mouse populations.

**Figure 6:**
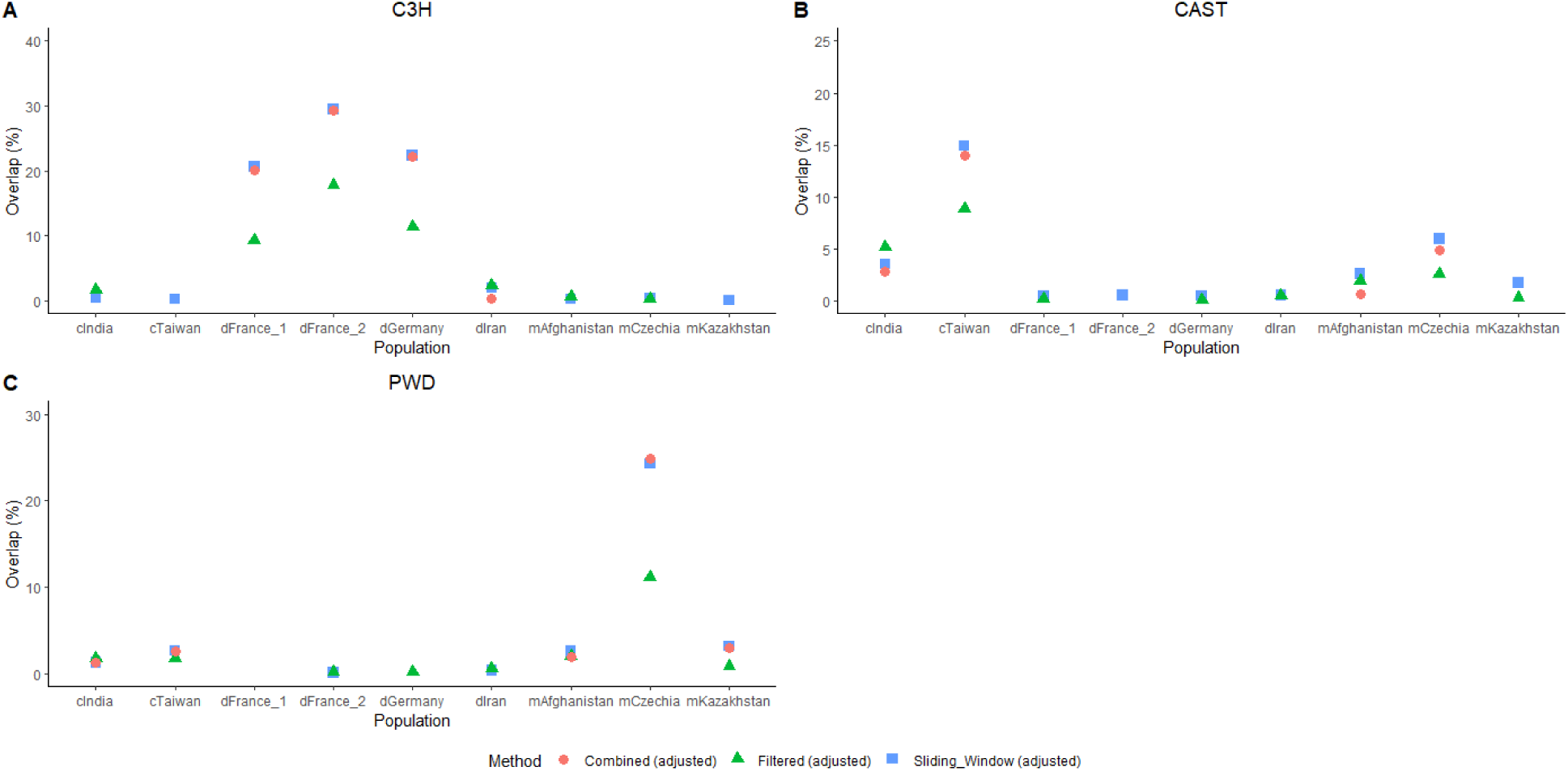
Wild mouse hotspots are more likely to overlap DSB hotspots of laboratory strains of the same sub-species. A) shows overlaps with C3H, a *M. m. domesticus* strain, B) shows overlap with CAST, a *M. m. castaneus* strain, C) shows overlaps with PWD, a *M. m. musculus* strain. All values are adjusted by subtracting the amount of overlap expected by chance (see Methods).

### Analysis of sex-specific recombination rate

LDHelmet yields sex-averaged recombination rate estimates, but, because the X chromosome only recombines in females, recombination rate comparisons between the X and autosomes stand to provide sex-specific information about female recombination.

Assuming wild mouse populations are at Hardy-Weinberg equilibrium, the mean *ρ*/bp of the X chromosome is expected to be 2/3rds the recombination rate of the autosomes, (as the X chromosome spends 2/3rds of its time in females). Remarkably, 8 of the 9 populations deviated from this expectation by more than 10% (Figure 7). In mice from cTaiwan, mCzechia, dGermany, dFrance_1, dFrance_2, and mKazakhstan, chrX recombination rates are higher than expected, suggesting that (i) overall recombination rates are elevated in females or (ii) that sex-specific demographic or selective histories have led to departures from HWE assumptions in these populations. Intriguingly, an opposite pattern is observed in the cIndia and dIran populations, with chrX recombination rates ( *ρ*/bp) falling below the expected value relative to the autosomes. Only the mAfghanistan population had a chrX recombination rate similar to the expectation (69%). These findings suggest that variation in the polarity of sex dimorphism for recombination rate may exist in wild mouse populations.

**Figure 7:**
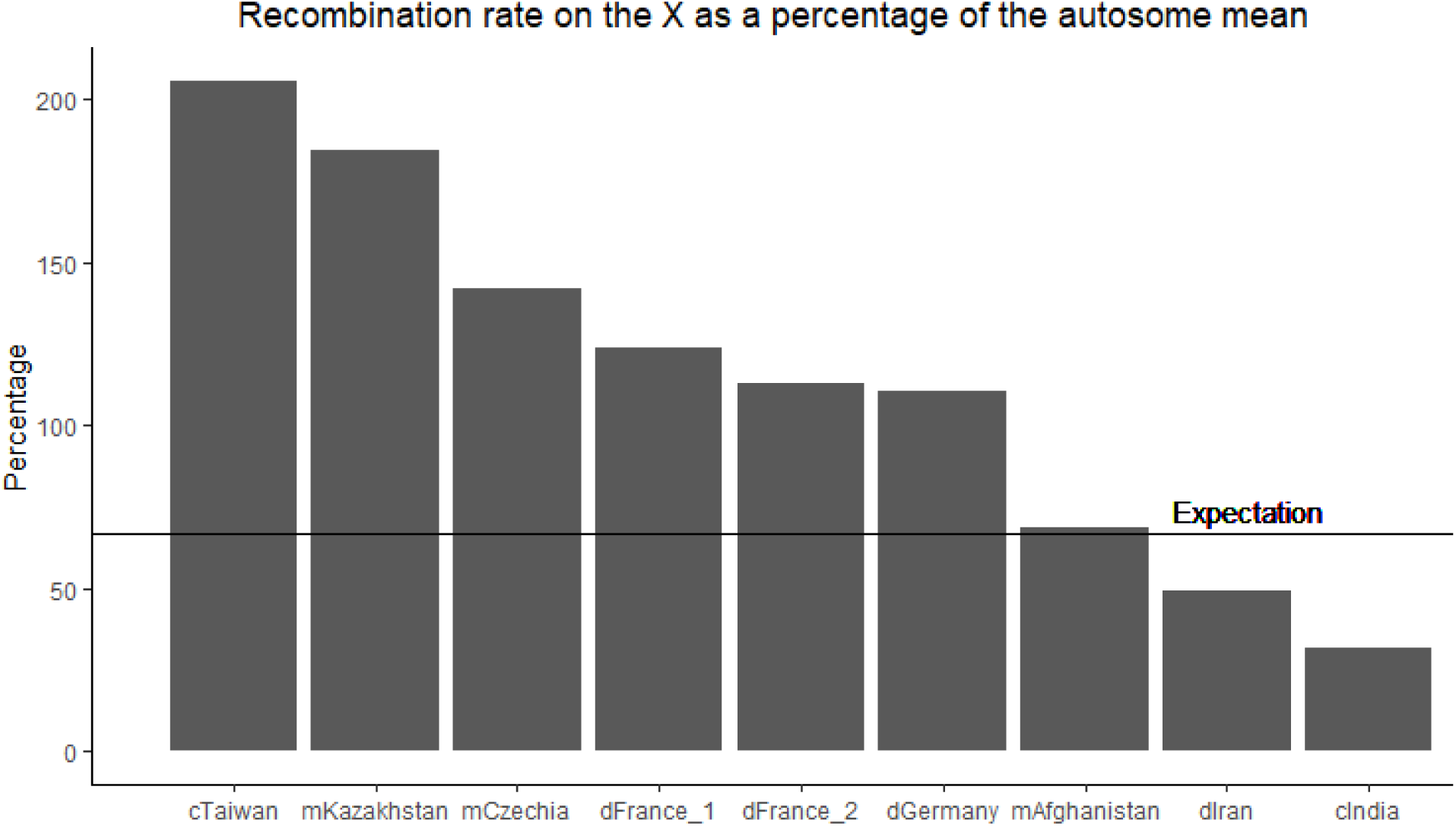
Different populations of wild house mice exhibit differences in scale and direction of sex-dimorphism in recombination rate. The recombination rate of the X chromosome was compared to the mean recombination rate of autosomes within each population and is expressed as a percentage of that mean. The expectation for the X chromosome (67%) is shown as a horizontal line.

### Impact of population evolutionary history on recombination rate estimation

LD-based methods for recombination rate estimation rest on a number of simplifying assumptions about population history that are rarely met in practice. Prior studies have demonstrated that non-random mating, changes in population size, the action of positive selection, and gene flow can lead to overestimation or underestimation of recombination rates. Model violations may also impact power to discover hotspots and lead to high rates of false positive hotspots (Reed and Tishkoff 2006; Zaitlen et al. 2017; Dapper and Payseur 2018; Samuk and Noor 2021). Each of the wild mouse populations examined here has experienced substantial changes in population size and bouts of positive selection in recent evolutionary history (Lawal et al. 2021).

To assess the impact of population demography on recombination rate estimation and hotspot discovery, we simulated genomic datasets for seven of the nine populations, invoking previously estimated demographic parameters for each population (see Methods) (Lawal et al. 2021). We then estimated recombination rates and inferred the positions of recombination hotspots in these simulated datasets. With the exception of cTaiwan, demographic history does not introduce overt biases in broad-scale *ρ* estimation; datasets simulated under population-specific demographic histories yield estimates of *ρ* that are not significantly skewed from the actual *ρ* value (Supp. File 8; Supp. Fig. 2). In the cTaiwan population, LDhelmet consistently underestimates *ρ* . This population experienced stronger historical bottlenecks than other wild mouse populations surveyed here, which likely accounts for this bias.

For the dGermany, mCzechia, cIndia, and dIran populations, power to detect hotspots exceeds 80%; for the remaining populations, power exceeds 50%. Importantly, false positives account for ≤10% of sliding window hotspots in all populations (Supp. Files 8-9). The false positive rate for filtered hotspots is notably higher, and hotspot detection power is slightly weaker using this method. Nonetheless, hotspots are localized with greater precision using this approach (Supp. File 9).

We conclude that, while differences in population history may contribute to population differences in the power to find hotspots, the demographic history of most wild mouse populations does not lead to strong skews in associated estimates or introduce large numbers of false positive hotspots.

## Discussion

For decades, inbred laboratory mice have been used as models to understand the molecular mechanisms and extent of variability in meiotic recombination. Indeed, studies in house mice helped lead to the initial discovery of *Prdm9* and its roles in hotspot specification (Parvanov et al. 2010). However, despite this progress, very little is known about how recombination landscapes vary or evolve in non-inbred, wild *Mus musculus*. Here, we used a population genomic approach to construct recombination maps for nine diverse populations of wild mice. Through extensive simulations, we demonstrate that our maps are robust to known departures from neutrality in these populations. Comparisons of both broad and fine-scale recombination rate divergence between populations and subspecies yielded several key findings.

First, we show that levels of genetic divergence between populations do not predict rates of broad-scale recombination rate divergence. Broad-scale map comparisons between populations of the same subspecies versus comparisons between populations derived from distinct *Mus musculus* subspecies yielded correlation values of similar magnitude. Our findings stand in contrast to predictions based on prior work. For example, Stevinson et al. found that correlations between broad-scale recombination maps decline with sequence divergence between great ape species (Stevison et al. 2016). The map correlations between house mouse populations are weaker than those reported between great ape species, even though *Mus musculus* subspecies diverged (∼0.3-0.5 MYA; ∼ 100,000 generations) more recently than humans and chimpanzees (∼6-8 MYA; 300,000 generations) (Geraldes et al. 2011; Langergraber et al. 2012; Amster and Sella 2016; Phifer-Rixey et al. 2020). Taken together, these findings suggest that the broad-scale recombination landscape evolves more quickly in house mice than in great apes, an outcome that may be attributable to taxon-specific differences in the functional constraints on the distribution of recombination events.

Second, we show that X chromosome recombination rates appear to evolve more slowly than autosomal recombination rates. In 35 of 36 population-level comparisons (See Supp. File 2), broad-scale recombination rates are more highly correlated along chrX than the autosomes, implying stricter conservation of the chrX recombination landscape. This finding aligns with prior work demonstrating that X chromosome recombination rates are more strongly correlated between cIndia and laboratory mice as compared to autosomes (Booker et al. 2017). This trend may be attributable, in part, to lower SNP density on the X. Indeed, we found that, in each of our nine surveyed populations, the X chromosome had the lowest SNP density (approximately 2-4x less than the autosomes of that population). However, the X chromosomes of cIndia and dIran have SNP densities similar to those of autosomes from other populations (1 SNP per 322 and 128 bp for dIran and cIndia, respectively), and the correlation between these two X chromosomes is still considerably higher than most autosome comparisons (Spearman’s *ρ* = 0.61 versus the autosome mean Spearman’s of all autosome comparisons *ρ* = 0.32). Additionally, there is a weak negative correlation between average SNP density between two populations and the strength of map correlation (See Supp. Fig. 3), indicating that lower SNP resolution is unlikely to provide a singular explanation for this observation.

Third, we uncover potential evidence of population differences in the magnitude and direction of sex dimorphism for recombination rate. Under a neutral Wright-Fisher model of evolution, mean X chromosome *ρ* is expected to equal 2/3 of the mean autosomal *ρ* estimate. Only the mAfghanistan population matches this neutral expectation (Figure 7). For most surveyed populations (cTaiwan, mKazakhstan, mCzechia, dGermany, and dFrance_1 and 2), the chrX *ρ* estimate exceeds neutral expectations based on the corresponding autosomal estimate. This result aligns with the common observation of higher global recombination rates in mouse females compared to males (Paigen et al. 2008; Dumont et al. 2009). However, the *ρ* estimates for chrX in the cIndia and dIran populations were less than expected based on the autosomal *ρ* estimates in these populations. Although higher female recombination rates present the dominant trend in inbred mouse genomes, cytogenetic investigations in inbred house mice have identified a select number of strains with higher male than female recombination rates (Peterson and Payseur 2021). Evidently, the polarity of sex dimorphism for global recombination rates can evolve rapidly. However, differences in demographic and selective history between males and females could bias X chromosome *ρ* estimates, leading to incorrect inferences about relative recombination rates between the sexes. Future work is needed to develop sex-specific models of evolutionary history for the populations investigated here and rigorously evaluate this potential interpretation. However, our results raise the possibility that the direction of the sex dimorphism for recombination rate varies between wild *Mus musculus* populations, and that previous observations of variation in the directionality of this dimorphism in inbred strains are not simply oddities of inbreeding.

In addition to these conceptual advances, we also developed a new method for the identification of hotspots in population data that fully utilizes the high density of SNPs in modern genome sequencing datasets. This ‘filtering’ method is simple to implement and detects hotspots at a finer resolution than the sliding window approach that has been used in prior studies (Booker et al. 2017; Shanfelter et al. 2019). Implementing this new method allowed for identification of an additional ∼11,000 new hotspots per population, and a total of more than 100,000 new hotspots for all nine populations combined. However, the filtering method failed to identify 115,978 hotspots called by the sliding window method, which suggests that both methods should be used in tandem to comprehensively identify hotspots in population data. The filtering method’s failure to detect these hotspots is potentially attributable to two reasons. First, the filtering method requires that a hotspot be comprised of at least three SNPs, while the sliding window method has no minimum SNP number requirements. In areas of the genome with lower SNP density, the sliding window method may be more likely to detect hotspots than the filtering method. Second, our implementation of the sliding window method required that hotspots be 10 times hotter than only the flanking 40kb regions, while the filtering method identified hotspots 10 times hotter than the mean of the entire chromosome. Differences in the recombination rate between the immediate flanking region and the entire chromosome undoubtedly allowed for some differential detection. Intriguingly, the mean *ρ*/bp of the filtered hotspots was on average nearly double the mean of the sliding window hotspots, and the same trend was also found when hotspots unique to the filtering method were compared to hotspots unique to the sliding window method. This indicates that the sliding window method misses a significant number of ‘very hot’ hotspots, likely because these hotspots are smaller than the sliding window.

We show that the number of detected hotspots per population scales with effective population size. This trend is expected if larger populations harbor greater *Prdm9* diversity, and thus a broader repertoire of recombination hotspot positions. Based on previous work, *M. m. musculus* is expected to have the smallest effective population size (*N_e_*= 100,000), followed by *M. m. domesticus* (160,000) (Salcedo et al. 2007), and with *M. m. castaneus* having the largest *N_e_* (580,000) (Geraldes et al. 2008). On average, about 30,000 hotspots were detected for *M. m. musculus*, 34,000 for *M. m. domesticus*, and 51,000 for *M. m. castaneus*. Within subspecies, hotspot numbers also varied between populations in a manner consistent with effective population size. The most dramatic example is a 1.6-fold difference in total number of hotspots detected between the India and Taiwan populations of *M. m. castaneus* (62,879 vs. 38,778 for cIndia and cTaiwan, respectively). This discrepancy again reflects known features of population history: the Taiwan population experienced a strong founding bottleneck that reduced its effective population size relative to ancestral populations of *M. m. castaneus* (Lawal et al. 2021). This bottleneck led to a genome-wide loss of diversity, including, presumably, a loss of allelic variation at the *Prdm9* locus, narrowing the suite of potentially active hotspot locations. Intriguingly though, this phenomenon of hotspot number scaling with population size is largely limited to hotspots detected by the filtering method, rather than the sliding window method. For the filtered hotspots, we detected on average 16,000, 20,000, and 43,000 hotspots for *M. m. musculus*, *M. m. domesticus*, and *M. m. castaneus*, respectively, while the sliding window method always detected an average of 24-25,500 hotspots per sub-species.

Hotspot location also varied greatly between populations, regardless of the hotspot identification method. These results extend prior observations of limited hotspot sharing between species (Stevison et al. 2016; Shanfelter et al. 2019) to the mouse model system. Remarkably, however, our work suggests that hotspot location varies greatly even between populations from the same *M. musculus* subspecies. This observation is at odds with significant hotspot sharing between human populations (Auton et al. 2012; Spence and Song 2019). Population genetic surveys of *Prdm9* allelic variation in wild-caught mice across the globe indicate an extensive number of *Prdm9* alleles segregating in nature (>100), with only 6 alleles shared between two subspecies and just 2 alleles shared between all three subspecies of *Mus musculus* (Buard et al. 2014). PRDM9 is known to interact epistatically with a locus on the X chromosome to cause hybrid male sterility in some intersubspecific experimental mouse crosses (Forejt et al. 2021). The entanglement of PRDM9 in a genetic incompatibility presumably restricts *Prdm9* gene flow in the wild and contributes to limited hotspot sharing between populations. Although the *Prdm9* genotype status of the individuals used to generate these LD recombination maps is not known and cannot be deciphered from short-read genome sequences, the lack of hotspot overlap between subspecies is consistent with high levels of population-private *Prdm9* allelic diversity in these wild mouse populations.

Although there is limited conservation of hotspots between wild populations, we observe appreciable levels of hotspot overlap between some wild mouse populations and hotspots ascertained by DMC1 ChIP-seq in laboratory inbred mice strains (Smagulova et al. 2016). In fact, sliding window hotspots in dGermany and dFrance overlapped more than 25% of DSB hotspots identified in C3H/He mice. A similar proportion of hotspot sharing was observed between DSB hotspots in PWD, a wild-derived inbred strain of *M. m. musculus* developed from wild-caught mice in the Czech Republic, and wild mice from the mCzechia population. These comparisons within subspecies retained >20% overlap after adjusting for the degree of overlap expected by chance. We note that, when we examined hotspots called by the filtering method in every population except cIndia, we found considerably less overlap with DSB hotspots (26,983 fewer overlaps, mean difference of 500 per comparison). This is likely because the sliding window hotspots are on average around three times longer in length than the filtered hotspots and are therefore more likely to overlap a DSB hotspot by chance. Indeed, our size-matched sliding window randomspots overlapped ∼11,000 more DSB hotspots on average than size-matched filtered randomspots. Our recombination maps effectively integrate over the historical *Prdm9* allelic diversity in each of our populations, but these trends suggest that several *Prdm9* alleles present in contemporary lab populations have left detectable footprints in the recombination landscape of wild mouse populations.

Overall, our findings expose remarkable divergence in the fine- and broad-scale recombination landscape between wild *Mus musculus* populations and subspecies. Evidently, the vast *Prdm9* allelic variation present in wild mouse populations has defined unique sets of genomic hotspots that have remained largely private to single populations for sufficiently long to render population-specific footprints in patterns of linkage disequilibrium. These results carry important practical implications for mouse genetics. Only a small subset of the *Prdm9* alleles found in wild mice are present in inbred mouse strains, a prospect that undoubtedly constrains mapping resolution in experimental crosses (and especially crosses between strains with identical *Prdm9* genotypes). Our fine-scale hotspot maps, combined with knowledge of the dominant *Prdm9* alleles in individual populations, stand to inform innovative experimental strategies for engineering diverse wild *Prdm9* alleles into laboratory strain genetic backgrounds. Such approaches could enable deliberate genetic manipulation of the crossover landscape and expedite efforts to fine map loci contributing to complex traits and disease.

## Materials and Methods

### Single nucleotide polymorphism data

We analyzed whole genome sequences from 99 wild *Mus musculus* (Davies 2015; Harr et al. 2016). These mice were trapped in nine different geographic locations on two continents. A basic summary of the data, including trapping location, sex, and subspecies identity, can be found in Table 1. This dataset features four populations of *Mus musculus domesticus*, three populations of *Mus musculus musculus*, and two populations of *Mus musculus castaneus*. Two of the *M. m. domesticus* populations sample mice from distinct locations in France; these populations were analyzed separately here and are designated as dFrance_1 (Harr et al. 2016) and dFrance_2 (Davies 2015).

Variants were called from whole genome sequences using the GATK best practices pipeline and GATK 4.1.8.1 (Van der Auwera and O’Connor 2020), as outlined in (Lawal et al. 2021). Single nucleotide polymorphisms (SNPs) were then filtered using a multistep process. First, the original VCF file containing all samples was split into nine files containing only samples and segregating sites from each population. Variants were then filtered using Vcftools 0.1.16 (Danecek et al. 2011). We retained biallelic sites with the Filter flag ‘PASS’, a minimum Quality score of 30, a minimum Genotype Quality score of 15, a minimum allele count of 2, and those that passed the Hardy-Weinberg equilibrium test (P > 0.0002). Additionally, SNPs were filtered based on the population’s mean read depth, and any sites with a read depth less than half or greater than double the population mean were excluded. This filter was applied to eliminate potential false positive calls due to read mismapping in structurally variable genomic regions.

### Estimating phase and switch-error rates

ShapeIt4 was used to infer haplotypes for each sample using standard parameters (Delaneau et al. 2019). To estimate the switch-error rate in our data, we paired phase-known X chromosomes from male samples to generated ‘pseudofemales’, as described (Booker et al. 2017). Briefly, reads mapping to the X chromosome from three to four males per population were merged to create all possible phase-known diploid combinations. Attempts to utilize only two males (therefore one pseudofemale) failed because ShapeIt4 requires multiple samples to infer phase. Only seven of the nine populations had sufficient male samples to be used for this analysis (mCzechia and cTaiwan had <3 males and could not be used). Variants were then called using GATK and filtered as described above. From each pseudofemale, we removed sites that were heterozygous in the true males (presumably SNPs located in the PAR), homozygous in the pseudofemale, or had missing data. After filtering, the pseudofemales were phased using ShapeIt4, and the resulting haplotypes converted into fasta format using bcftools (v 1.9.1) *consensus* and the mm10 reference sequence (Danecek et al. 2021). These whole chromosome fasta sequences were then pared down to include only sites segregating in the pseudofemale. The inferred haplotypes from a pseudofemale were next compared to the phase-known sequences of the two donor male chrX sequences. The switch-error rate was defined as the number of switch-errors that occurred, divided by the total number of opportunities for a switch to occur (*i.e*., the total number of SNPs minus 1).

### LD-based recombination map construction

LDHelmet v1.10 was used to estimate the population scaled recombination rate for each chromosome in each of the nine *M. musculus* populations (Chan et al. 2012). Parameters were set based on developer recommendations and previously published work (Chan et al. 2012; Booker et al. 2017), with a few modifications. Briefly, before running the rjmcmc, haplotype configuration files were generated using a window size of 50. Likelihood lookup tables were constructed across a grid of population scaled recombination rates (0.0 0.1 10.0 1.0 100.0) and using subspecies-specific population mutation rates, assuming a common genomic mutation rate of 0.5 x 10^-8^ bp/generation (Uchimura et al. 2015) and effective population sizes of 160,000, 580,000, and 100,000 for *domesticus*, *castaneus*, and *musculus*, respectively (Salcedo et al. 2007; Geraldes et al. 2008). To improve accuracy of sampling, we computed 11 Pade coefficients using the same population-scaled mutation rate estimates. Once these preparatory files were generated, the rjmcmc was run using a window size of 50, a subspecies-specific mutation matrix, ancestral priors (see below), a partition length of 50,000 SNPs, and either a block penalty of 100 (broad-scale map) or 10 (fine-scale map). The rjmcmc program was then run for 1,000,000 iterations for each block penalty, with the first 100,000 iterations discarded as burn-in

Ancestral priors were calculated using *M. caroli*, *M. spretus* and *M. pahari*, where alleles matching all three species, or matching in two but missing in the third, were considered the ancestral allele. To account for potential allele misspecification, the presumed ancestral allele was assigned a weight of 0.91, and the other three possible states were assigned a weight of 0.03. If the ancestral allele state could not be inferred, the overall frequency of that particular nucleotide in the mm10 reference genome was used.

### Conversion between population-scaled and genetic map distance

LDHelmet outputs estimates of recombination between adjacent SNPs in *ρ*/bp units. To convert this quantity into more readily interpretable cM/Mb units, we first summed the *ρ*/bp estimates across each chromosome to determine the total population-scaled recombination rate. For each pair of adjacent SNPs on the map, we then calculated the proportional contribution to total *ρ*. This percentage was then multiplied by the length of each chromosome in cM units, as estimated from the current gold-standard mouse genetic linkage map (Cox et al. 2009).

### Map comparisons

Spearman’s correlation was used to assess similarity of the recombination distribution (in terms of cM/Mb) between each wild mouse population. Correlations were assessed for whole genome comparisons, as well as for individual chromosomes. To gauge the strength of the correlation between two maps that could be expected due to chance, we generated 100 random permutations of *ρ* estimates in 1 Mb segments across each population’s genome. An empirical *P*-value was estimated as the fraction of simulated comparisons in excess of the observed Spearman’s *ρ*-statistic.

A prior study used different methodology to create LD-based recombination maps for the cIndia population studied here (Booker et al. 2017). To compare our cIndia maps to the prior map for this population, SNP positions on this earlier map were converted from the mm9 to mm10 coordinate system using LiftOver from the UCSC tool suite (Hinrichs et al. 2006).

### Identification of hotspots

The fine-scale recombination map from each population was used to identify putative recombination hotspots using two approaches. We first identified hotspots using a conventional ‘sliding-window’ approach (Shanfelter et al. 2019), with minor modifications. In brief, the mean *ρ* of each 1 kb window (0.5kb slide) was compared to the mean *ρ* of the flanking 40 kb regions. If *ρ* in the 1 kb target segment was greater than 10 times the population-scaled recombination rate of the flanking regions, the region was determined to be a hotspot.

This sliding window method does not fully take advantage of the high SNP density in modern datasets (here, 1 SNP every ∼60-300 bp). To fully leverage the high SNP density in our dataset, we developed and implemented a new method for hotspot detection. Briefly, a segment of DNA between adjacent SNPs was labeled a putative hotspot if its *ρ*/bp was at least 10x the chromosome-wide mean *ρ*. Putative hotspots with shared SNPs were then merged into a single candidate hotspot. Only candidate hotspots with >2 SNPs and <5 kb in length were retained. We set a minimum requirement of 3 SNPs contained in a hotspot to reduce the risk of false-positive hotspots due to genotyping or haplotype switch errors. A maximum hotspot length of 5 kb was invoked based on extensive prior estimates of likely hotspot size (Paigen et al. 2008; Altshuler et al. 2010; Tsai et al. 2010).

This new method, which we term the ‘filtering’ approach, yielded some pairs of adjacent hotspots separated by only 2 SNPs. These cases may reflect two independent, closely positioned hotspots, but it is also plausible the two hotspots are actually a single hotspot that was erroneously split in two, potentially due to genotyping error. We took a conservative approach and merged any hotspots separated by 2 SNPs and that were ≤ 1 kb apart. Hotspots separated by 2 SNPs and positioned >1 kb apart were retained as independent hotspots. Hotspots separated by 3 or more ‘cold’ SNPs were always treated as individual hotspots.

Bedtools intersect (v2.29.2) was used to identify hotspots detected by both the ‘sliding-window’ and ‘filtering’ approaches, imposing a minimum overlap requirement of 1 bp. Overlapping hotspots between the calling methods were then merged and combined with hotspots unique to each method to create a total set of hotspots for each population of mice.

### Identification of coldspots

We used a method similar to the filtering hotspot approach outlined above to identify coldspots, or areas of comparatively low recombination. Specifically, a segment was inferred to be a coldspot if *ρ*/bp was less than 1/10th the chromosome average and if it contained at least 3 SNPs. No minimum or maximum length requirements were imposed on coldspots.

### Generation of ‘randomspots’

To assess various outcomes expected by chance, we generated 100 sets of random, size-matched genomic segments to mimic both the filtered and sliding window hotspots detected on each chromosome in each population using a custom Python script (Supp. File 10). We refer to these simulated regions as ‘randomspots.’

### Comparison of hotspots between populations

We analyzed each population-specific set of hotspots (filtered or sliding window) for overlap between populations within each subspecies, as well as across subspecies. When comparing populations within a subspecies, bedtools intersect was used to find hotspots with at least 1 bp of overlap, at least 50% overlap (-f 0.5 and -F 0.5 -e; partial overlap), or 100% overlap (-f 1.0 -F 1.0-e; complete overlap). When comparing across subspecies, only hotspots with at least 50% overlap were examined (-f 0.5 and -F 0.5 -e).

### Characterizing the genomic distribution of hotspots

We analyzed our hotspots for proximity to transcription start sites (TSS) and repeat elements. Bedtools intersect was used to find hotspots overlapping at least 1 bp of an annotated TSS (refTSS) or repetitive element (repeatmasker) (Smith et al. 2013; Abugessaisa et al. 2019). Bedtools closest was used to find the closest hotspots to each TSS, along with the distance between them. Fisher’s Exact tests were used to identify repetitive elements with differential enrichment between hot- and cold-spots. The percent repeat composition (in terms of bp) of hotspots and coldspots was compared using Welch’s t-test. The effect of hotspot size on the percent composition of repeats was analyzed by beta regression. Due to the high number of extreme values (0.0 and 1.0, indicating 0 or 100% repeat composition) in this dataset, repeat fractions were first transformed using the formula (y * (n-1) + 0.5)/n, where n is the sample size and y is the repeat proportion (Smithson and Verkuilen 2006).

We also analyzed our hotspots for overlap with previously published double-strand break (DSB) hotspots ascertained using ChIP-seq against DMC1, a protein that binds to the ends of DNA DSB breaks (Smagulova et al. 2016). Overlap was assessed using bedtools intersect, with a requirement for at least 1 bp overlap.

### Motif enrichment

For each population, hotspots were compared to a random subset of 50,000 coldspots to identify hotspot-enriched motifs. Alleles present at a frequency of greater than 50% frequency in the focal population were used in the population-specific consensus hotspot sequence, with lower frequency alleles replaced by the reference nucleotide. We then used STREME (v5.3.3) to identify a maximum of 10 hotspot-enriched motifs 8-15 bp in length per population (Bailey 2021). Tomtom (v5.4.1) was used to find known motifs in the HOCOMOCO v11 Full database that exhibited high sequence similarity to our hotspot-enriched sequences (Gupta et al. 2007).

### Coalescent simulations

Prior work has established that historical bottlenecks and population expansions can leave footprints in population-wide patterns of LD that manifest as false-positive hotspots (Dapper and Payseur 2018). We conducted a series of coalescent simulations to assess the performance of LDHelmet in the face of departures from neutral demographic assumptions. Briefly, we used msHOT to simulate 100kb haplotypes according to each population’s demographic history (Hellenthal and Stephens 2007). The number of simulated haplotypes was specified to match the number of samples for each population. Effective population sizes and population specific demographic parameters were previously estimated for seven of the nine wild mouse populations profiled here, with the exception of the individual French populations (Lawal et al. 2021). We invoke these previously estimated values to generate haplotypes under models that reflect each population’s unique evolutionary history. For all simulations, we assume a background recombination rate of 0.05 cM/Mb, corresponding to *ρ* = 0.02/kb. A single 2-kb hotspot with recombination rate 100x the background was simulated in the center of each 100kb region. For each population, we performed 100 replicate simulations. The executed commands are provided in Supp. File 9.

For comparison, we also simulated data for a single representative population under a neutral demographic model with no historical population contraction or expansion. As above, we simulated a single 2-kb hotspot centered within a 100kb region and invoke the same recombination rate assumptions. We assume an effective population size of 100,000 – comparable to the magnitude of many of the mouse populations used in this investigation – and 8 samples (16 haplotypes).

The simulation output from msHOT was then converted to fasta format using a custom R script. Sequences were simulated from the observed nucleotide frequencies in the mm10 reference genome. Derived alleles were sampled using the mutation probabilities specified by the *M. m. domesticus* transition matrix (described above). While all simulations rely on a single transition matrix for convenience, we note that there are negligible differences in observed mutation transition probabilities across subspecies. At each mutation site, the ancestral allele was assigned a weight of 0.91, with the other three possible states assigned a weight of 0.03, as above. Fasta files were used as input into LDhelmet. Hotspot discovery was performed as outlined above. Due to the small size of simulated fragments *ρ*, was only estimated using a block penalty of 10, providing greater resolution for detection of recombination rate heterogeneity.

We estimated the power to detect simulated hotspots and the rate of false positive discovery for each population. Hotspots detected within 10kb of the start or end of simulated sequence were excluded to minimize the contribution of artifacts due to edge effects. Power was calculated as the proportion of simulation replicates for which the midpoint of a detected hotspot was <5 kb from the simulated hotspot position. The false positive rate was computed as the fraction of simulations with ≥ 1 detected hotspot outside this 5kb window.

## Acknowledgements

This work was supported by a National Science Foundation CAREER award to BLD (DEB 1942620). We thank Drs. Michael White and Alice Shanfelter for sharing a Perl script for implementing the sliding window approach for hotspot identification. We also thank members of the Dumont Laboratory for feedback on this work. In particular, we thank Dr. Raman Lawal for providing the variant calls for these samples and Alexis Garretson for guidance on data presentation.

## Data Availability

The data underlying this article are available in the article and in its online supplementary material. Custom scripts are available from the authors upon request.

