## Supplemental Figure 1 for "Rapid evolution of the fine-scale recombination landscape in wild house mouse (*Mus musculus*) populations"

| Population | Motif | Image | Abundance (%) |
|---|---|---|---|
| **dFrance_1** | GCAGCAKCV | 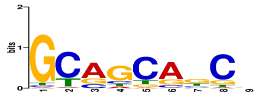 | 95.48 |
| **dFrance_2** | GAWGCTRCTRCC | 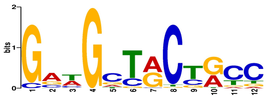 | 24.36 |
| **dFrance_2** | AAGGTGGCA | 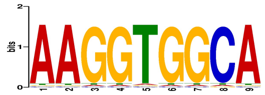 | 0.06 |
| **dFrance_2** | GCAGCAGCC | 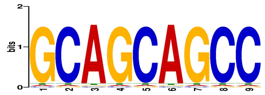 | 0.04 |
| **dGermany** | SWGCWGCWS | 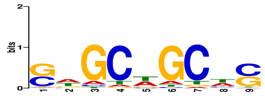 | 77.18 |
| **dIran** | CKGCAGCRGC | 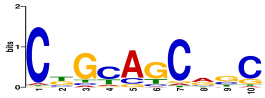 | 50.79 |
| **dIran** | WRVWSCWG | 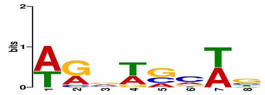 | 30.54 |
| **dIran** | SAGAAACTGC | 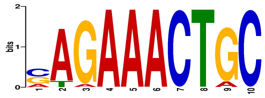 | 0.05 |
| **mAfghanistan** | None detected | | - |
| **mCzechia** | GSYTRYARSC | 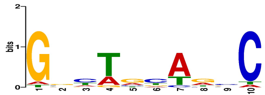 | 87.83 |
| **mKazakhstan** | AATGGTGCTGGCA | 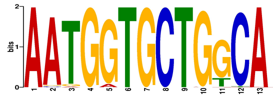 | 0.99 |
| **mKazakhstan** | AGATTTTGCTG | 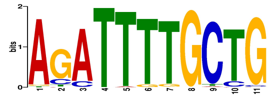 | 2.20 |
| **mKazakhstan** | AGAAGCTGTTAGMW | 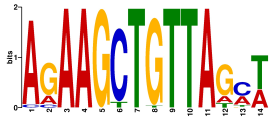 | 0.29 |
| **cIndia** | CAGCAGCAC | 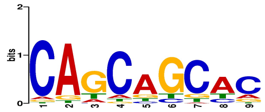 | 21.56 |
| **cIndia** | AAGAAGGC | 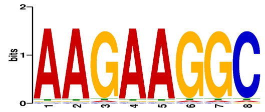 | 0.23 |
| **cTaiwan** | AARARGSW | 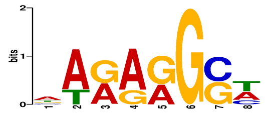 | 4.66 |

| Population | Motif | Image | Abundance (%) |
|---|---|---|---|
| **dFrance_1** | CAGAAGCA | 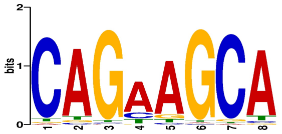 | 97.70 |
| **dFrance_2** | None detected | | - |
| **dGermany** | AASRWMNHVSNRV | 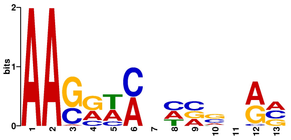 | 97.54 |
| **dGermany** | GCHRCAGBHHB | 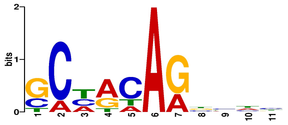 | 97.01 |
| **dGermany** | AHVMTGMWNNM | 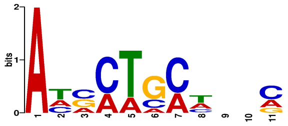 | 96.37 |
| **dIran** | RDGRCWGB | 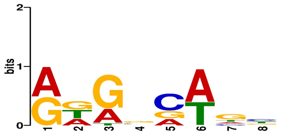 | 72.71 |
| **mAfghanistan** | None detected | | - |
| **mCzechia** | AAGAAGCTG | 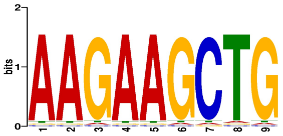 | 0.04 |
| **mKazakhstan** | CTGTAGCA | 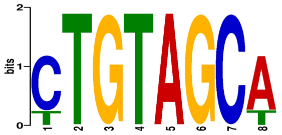 | 0.13 |
| **mKazakhstan** | TGGCACCAG | 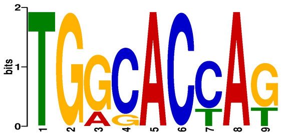 | 0.07 |
| **cIndia** | None detected | | - |
| **cTaiwan** | None detected | | - |
