## Supplementary figures and images for "Rapid evolution of the fine-scale recombination landscape in wild house mouse (*Mus musculus*) populations"

### Supplemental Figure 2

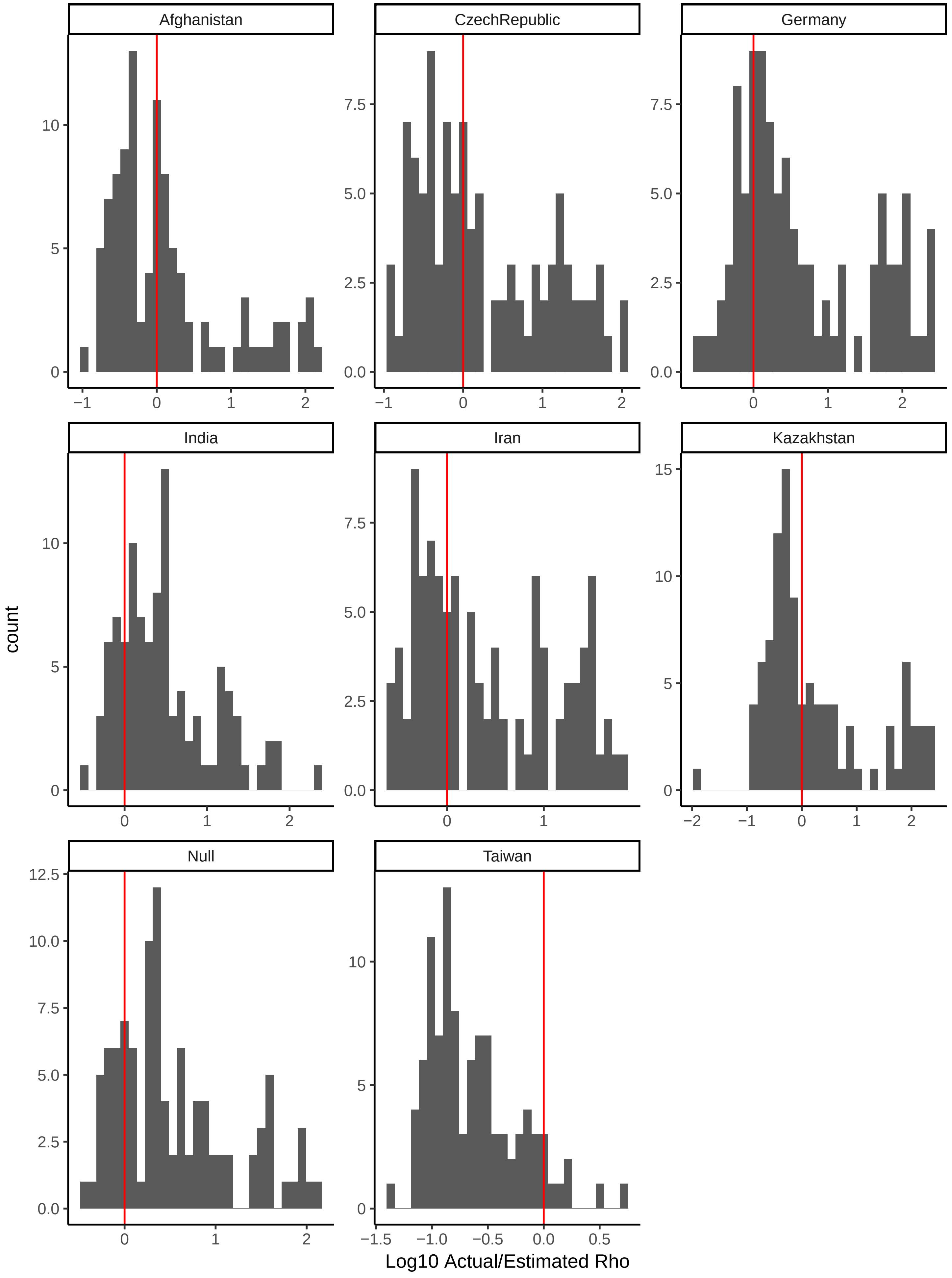

### Supplemental Figure 3

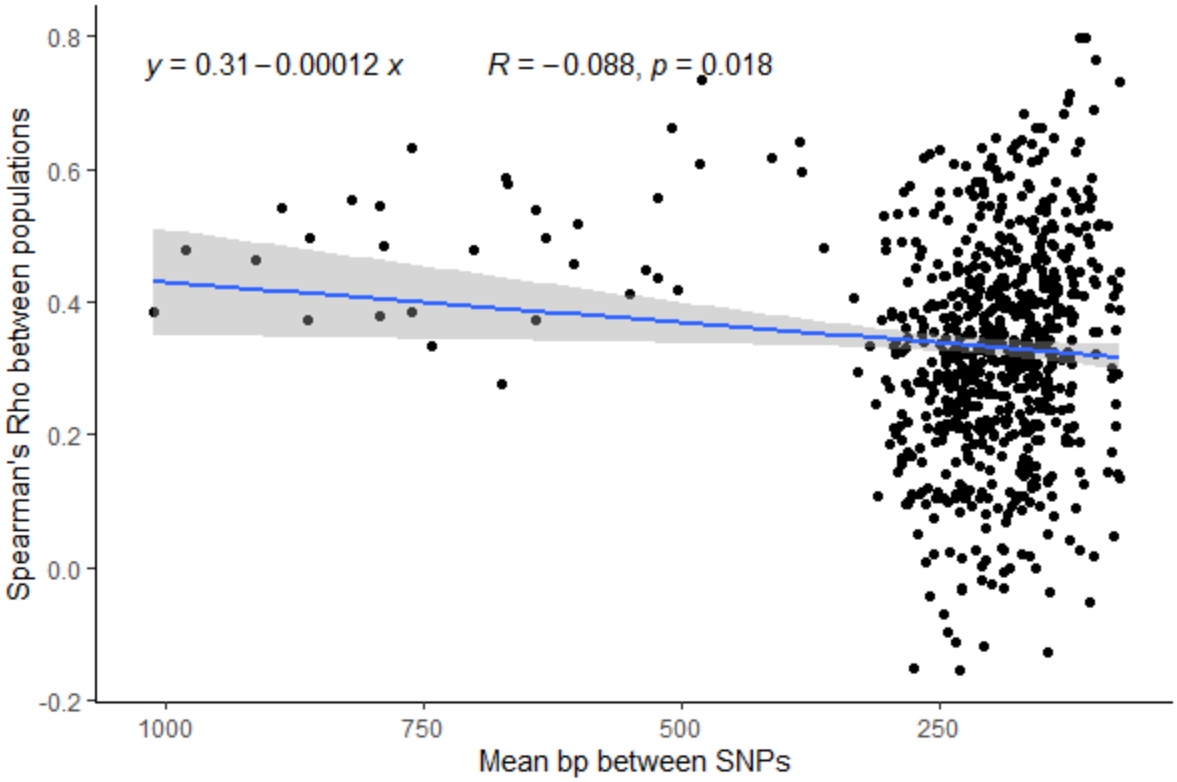
