## Supplemental File 3 for "Rapid evolution of the fine-scale recombination landscape in wild house mouse (*Mus musculus*) populations"

| | Total | Percentage (%) |
|---|---|---|
| Raw Potential Hotspots | 2,586,882 | |
| 2 SNP Hotspots | 2,363,177 | 91.35 |
| >5 kb | 4,800 | 0.19 |
| 2 cold SNPS in between, <1 kb separation | 4,048 | 0.16 |
| 2 cold SNPS in between, >1 kb separation | 140 | 0.005 |
| Final, Filtered Hotspots | 214,717 | 8.30 |
