## Supplemental File 5 for "Rapid evolution of the fine-scale recombination landscape in wild house mouse (*Mus musculus*) populations"

| | Percent bp are from a repeat element (%) | | |
|---|---|---|---|
| | **Coldspots** | **Sliding Window Hotspots** | **Filtered Hotspots** |
| **mAfghanistan** | 44.5* | 43.3** | 44.3* |
| **mCzech** | 42.3* | 42.8** | 48.4*** |
| **mKazakhstan** | 51.2* | 43.2** | 58.4*** |
| **dIran** | 44.0* | 42.1** | 42.1** |
| **dGermany** | 44.5* | 41.0** | 45.3*** |
| **dFrance_1** | 42.8* | 42.4* | 47.2** |
| **dFrance_2** | 43.5* | 41.1** | 45.2*** |
| **cTaiwan** | 43.5* | 46.7** | 51.2*** |
| **cIndia** | 50* | 40** | 38.3*** |
