## Supplemental File 6 for "Rapid evolution of the fine-scale recombination landscape in wild house mouse (*Mus musculus*) populations"

| Strain | Wild Population | # DSB Hotspots | # Overlaps | Random Overlaps | # Overlaps – Corrected | % Overlap - Corrected |
|---|---|---|---|---|---|---|
| 13R | mAfghanistan | 14,752 | 737 | 636 | 101 | 0.68 |
| | mCzechia | | 496 | 453 | 43 | 0.29 |
| | mKazakhstan | | 516 | 517 | -1 | -0.01 |
| | dIran | | 945 | 695 | 250 | 1.69 |
| | dGermany | | 722 | 439 | 283 | 1.92 |
| | dFrance_1 | | 865 | 488 | 377 | 2.56 |
| | dFrance_2 | | 897 | 497 | 400 | 2.71 |
| | cTaiwan | | 460 | 446 | 14 | 0.09 |
| | cIndia | | 564 | 534 | 30 | 0.20 |
| B6 | mAfghanistan | 19,528 | 960 | 828 | 132 | 0.68 |
| | mCzechia | | 705 | 586 | 119 | 0.61 |
| | mKazakhstan | | 603 | 673 | -70 | -0.36 |
| | dIran | | 1501 | 891 | 610 | 3.12 |
| | dGermany | | 3805 | 568 | 3237 | 16.58 |
| | dFrance_1 | | 2350 | 631 | 1719 | 8.80 |
| | dFrance_2 | | 2924 | 645 | 2279 | 11.67 |
| | cTaiwan | | 594 | 578 | 16 | 0.08 |
| | cIndia | | 746 | 690 | 56 | 0.29 |
| C3H | mAfghanistan | 14,645 | 764 | 727 | 37 | 0.25 |
| | mCzechia | | 580 | 519 | 61 | 0.42 |
| | mKazakhstan | | 612 | 602 | 10 | 0.07 |
| | dIran | | 1093 | 802 | 291 | 1.99 |
| | dGermany | | 3779 | 504 | 3275 | 22.36 |
| | dFrance_1 | | 3582 | 562 | 3020 | 20.62 |
| | dFrance_2 | | 4889 | 575 | 4314 | 29.46 |
| | cTaiwan | | 549 | 520 | 29 | 0.20 |
| | cIndia | | 689 | 626 | 63 | 0.43 |
| CAST | mAfghanistan | 15,068 | 1063 | 676 | 387 | 2.57 |
| | mCzechia | | 1371 | 476 | 895 | 5.94 |
| | mKazakhstan | | 799 | 542 | 257 | 1.71 |
| | dIran | | 807 | 733 | 74 | 0.49 |
| | dGermany | | 539 | 466 | 73 | 0.48 |
| | dFrance_1 | | 590 | 517 | 73 | 0.48 |
| | dFrance_2 | | 602 | 527 | 75 | 0.50 |
| | cTaiwan | | 2718 | 473 | 2245 | 14.90 |
| | cIndia | | 1089 | 565 | 524 | 3.48 |
| MOL | mAfghanistan | 15,768 | 1214 | 660 | 554 | 3.51 |
| | mCzechia | | 1467 | 473 | 994 | 6.30 |
| | mKazakhstan | | 1084 | 544 | 540 | 3.42 |
| | dIran | | 991 | 722 | 269 | 1.71 |
| | dGermany | | 605 | 458 | 147 | 0.93 |
| | dFrance_1 | | 611 | 505 | 106 | 0.67 |
| | dFrance_2 | | 697 | 525 | 172 | 1.09 |
| | cTaiwan | | 689 | 468 | 221 | 1.40 |
| | cIndia | | 855 | 566 | 289 | 1.83 |
| PWD | mAfghanistan | 14,539 | 1123 | 744 | 379 | 2.61 |
| | mCzechia | | 4066 | 535 | 3531 | 24.29 |
| | mKazakhstan | | 1077 | 613 | 464 | 3.19 |
| | dIran | | 878 | 821 | 57 | 0.39 |
| | dGermany | | 512 | 521 | -9 | -0.06 |
| | dFrance_1 | | 551 | 578 | -27 | -0.19 |
| | dFrance_2 | | 612 | 597 | 15 | 0.10 |
| | cTaiwan | | 919 | 533 | 386 | 2.65 |
| | cIndia | | 832 | 645 | 187 | 1.29 |
