## Supplemental File 7 for "Rapid evolution of the fine-scale recombination landscape in wild house mouse (*Mus musculus*) populations"

| Strain | Wild Population | # DSB Hotspots | # Overlaps | Random Overlaps | # Overlaps – Corrected | % Overlap - Corrected |
|---|---|---|---|---|---|---|
| 13R | mAfghanistan | 14,752 | 330 | 332 | -99 | -0.67 |
| | mCzechia | | 187 | 177 | -43 | -0.29 |
| | mKazakhstan | | 152 | 235 | -150 | -1.02 |
| | dIran | | 670 | 451 | 95 | 0.64 |
| | dGermany | | 280 | 215 | 4 | 0.03 |
| | dFrance_1 | | 322 | 202 | 64 | 0.43 |
| | dFrance_2 | | 450 | 291 | 82 | 0.56 |
| | cTaiwan | | 280 | 386 | -213 | -1.44 |
| | cIndia | | 843 | 713 | -60 | -0.41 |
| B6 | mAfghanistan | 19,528 | 489 | 429 | 87 | 0.45 |
| | mCzechia | | 220 | 230 | -1 | -0.01 |
| | mKazakhstan | | 209 | 302 | -85 | -0.44 |
| | dIran | | 1089 | 575 | 532 | 2.72 |
| | dGermany | | 1805 | 276 | 1540 | 7.89 |
| | dFrance_1 | | 954 | 258 | 702 | 3.59 |
| | dFrance_2 | | 1571 | 368 | 1215 | 6.22 |
| | cTaiwan | | 374 | 493 | -107 | -0.55 |
| | cIndia | | 1095 | 903 | 204 | 1.04 |
| C3H | mAfghanistan | 14,645 | 426 | 402 | 94 | 0.64 |
| | mCzechia | | 219 | 221 | 42 | 0.29 |
| | mKazakhstan | | 232 | 294 | -3 | -0.02 |
| | dIran | | 801 | 557 | 350 | 2.39 |
| | dGermany | | 1882 | 265 | 1667 | 11.38 |
| | dFrance_1 | | 1565 | 252 | 1363 | 9.31 |
| | dFrance_2 | | 2903 | 356 | 2612 | 17.84 |
| | cTaiwan | | 381 | 481 | -5 | -0.03 |
| | cIndia | | 950 | 891 | 237 | 1.62 |
| CAST | mAfghanistan | 15,068 | 641 | 358 | 283 | 1.88 |
| | mCzechia | | 581 | 196 | 385 | 2.56 |
| | mKazakhstan | | 285 | 252 | 33 | 0.22 |
| | dIran | | 551 | 483 | 68 | 0.45 |
| | dGermany | | 240 | 229 | 11 | 0.07 |
| | dFrance_1 | | 233 | 217 | 16 | 0.11 |
| | dFrance_2 | | 295 | 311 | -16 | -0.11 |
| | cTaiwan | | 1752 | 414 | 1338 | 8.88 |
| | cIndia | | 1537 | 766 | 771 | 5.12 |
| MOL | mAfghanistan | 15,768 | 711 | 355 | 356 | 2.26 |
| | mCzechia | | 584 | 193 | 391 | 2.48 |
| | mKazakhstan | | 432 | 253 | 179 | 1.14 |
| | dIran | | 658 | 484 | 174 | 1.10 |
| | dGermany | | 293 | 226 | 67 | 0.42 |
| | dFrance_1 | | 224 | 214 | 10 | 0.06 |
| | dFrance_2 | | 367 | 309 | 58 | 0.37 |
| | cTaiwan | | 489 | 419 | 70 | 0.44 |
| | cIndia | | 1300 | 766 | 534 | 3.39 |
| PWD | mAfghanistan | 14,539 | 710 | 408 | 302 | 2.08 |
| | mCzechia | | 1851 | 225 | 1626 | 11.18 |
| | mKazakhstan | | 419 | 299 | 120 | 0.83 |
| | dIran | | 653 | 568 | 85 | 0.58 |
| | dGermany | | 288 | 266 | 22 | 0.15 |
| | dFrance_1 | | 234 | 252 | -18 | -0.12 |
| | dFrance_2 | | 399 | 361 | 38 | 0.26 |
| | cTaiwan | | 738 | 485 | 253 | 1.74 |
| | cIndia | | 1161 | 901 | 260 | 1.79 |

Filtered
